# A brain circuit and neuronal mechanism for decoding and adapting to change in daylength

**DOI:** 10.1101/2023.09.11.557218

**Authors:** G Maddaloni, YJ Chang, RA Senft, SM Dymecki

## Abstract

Changes in daylight amount (photoperiod) drive pronounced alterations in physiology and behaviour^1,2^. Adaptive responses to seasonal photoperiods are vital to all organisms – dysregulation is associated with disease, from affective disorders^3^ to metabolic syndromes^4^. Circadian rhythm circuitry has been implicated^5,6^ yet little is known about the precise neural and cellular substrates that underlie phase synchronization to photoperiod change. Here we present a previously unknown brain circuit and novel system of axon branch-specific and reversible neurotransmitter deployment that together prove critical for behavioural and sleep adaptation to photoperiod change. We found that the recently defined neuron type called mr*En1-Pet1*^7^ located in the mouse brainstem Median Raphe Nucleus (MRN) segregates serotonin versus VGLUT3 (here proxy for the neurotransmitter glutamate) to different axonal branches innervating specific brain regions involved in circadian rhythm and sleep/wake timing^8,9^. We found that whether measured during the light or dark phase of the day this branch-specific neurotransmitter deployment in mr*En1-Pet1* neurons was indistinguishable; however, it strikingly reorganizes on photoperiod change. Specifically, axonal boutons but not cell soma show a shift in neurochemical phenotype upon change away from equinox light/dark conditions that reverses upon return to equinox. When we genetically disabled the deployment of VGLUT3 in mr*En1-Pet1* neurons, we found that sleep/wake periods and voluntary activity failed to synchronize to the new photoperiod or was significantly delayed. Combining intersectional rabies virus tracing and projection-specific neuronal silencing *in vivo*, we delineated a Preoptic Area-to-mr*En1Pet1* connection responsible for decoding the photoperiodic inputs, driving the neurochemical shift and promoting behavioural synchronization. Our results reveal a previously unrecognized brain circuit along with a novel form of periodic, branch-specific neurotransmitter deployment that together regulate organismal adaptation to photoperiod changes.

## Main

Sunlight potently influences life across all phylogenetic kingdoms. Synchronized to the 24-hour light/dark cycle are biological processes fundamental to energy efficiency, cellular and organismal homeostasis, and ultimately survival^9^. Animals have evolved exquisite mechanisms – collectively known as circadian rhythms orchestrated by the hypothalamic suprachiasmatic nucleus (SCN) – that enable coordinated anticipation of alternating day/night and dictate the temporal order of physiology and behaviour including metabolism, hormone secretion, physical activity, and sleep/wakefulness^9,10^. Equally vital are adaptive responses to changes in the duration of daily light exposure (photoperiod)^1,11^ such as those that occur with time of year especially at high latitudes or with trans meridian travel, shift work, or screen time. Compromised ability to phase shift to new photoperiods has deleterious effects on human health and can cause circadian and sleep-timing abnormalities that are tightly linked to affective^2,3^, neurodegenerative^12^, cardiovascular^13^, as well as metabolic disorders^4^, severely impacting quality of life. Changes in photoperiod duration and timing can induce switching of neurotransmitter usage by certain neurons in the SCN^6^ and their post-synaptic targets in the Paraventricular Nucleus of the Hypothalamus^14^, thus implicating the core circadian rhythm circuitry^5^ in phase adaptation to changes in light exposure. Little is known, however, about the precise causative mechanisms underlying phase synchronization at the molecular and circuit level.

Here we report the involvement of a subgroup of Median Raphe (mr) Nucleus (MRN) serotonin (5-hydroxytryptamine, 5-HT) neurons and associated brain circuits in adaptation to photoperiod change. MRN neurons provide the sole and dense 5-HTergic input to the SCN^15^ and potently modulate circadian rhythms^16–19^. Additionally, photoperiod changes have been associated with transcriptional^20^, electrophysiological^21^, and structural modifications^22^ in some 5-HTergic neurons, thus suggesting a potential yet unexplored role in modulating synchronization to day-length changes. Combining intersectional genetics, viral circuit tracing, and pathway-specific genetic and neuronal manipulations with measurements of physical and sleep/wake activity, we discovered in mice a preoptic area-to-brainstem-to-hypothalamus neural circuit and novel phenomenon of axon branch-specific and periodic neurotransmitter deployment that together prove critical for behavioural and sleep adaptation to seasonally relevant photoperiod change. Thus, identified is a cellular mechanism as well as a circuit that links internal rhythms to external stimuli.

### Brain region-specific and differential neurotransmitter deployment by axon collaterals of the dual serotonin-glutamate neuron type mr*En1-Pet1*

Previously, we identified mr*En1-Pet1* neurons as a molecularly and developmentally distinct subtype of 5-HT-producing neuron resident in the mouse MRN^7,23,24^ (Fig. 1a-c). As indicated by the name mr*En1-Pet1*, these median raphe (mr) neurons can be accessed genetically through the combined expression of the gene *En1* encoding a fate-specifying homeobox transcription factor, and *Pet1* (aka *Fev)* encoding an ETS-family transcriptional regulator of serotonergic pathway genes. To visualize and manipulate these neurons, we use mice transgenic for *Pet1-Flpe*^23^ and *En1-cre*^*51*^ along with dual Cre/Flp-dependent (intersectional) reporters (e.g. *RC-FrePe*^25^, *RC-FPSit*^26^; Fig. 1b) or effector transgenes^27^, or engineered stereotaxically injected viruses that allow for spatial targeting of just the mr (as opposed to dorsal raphe) group of *En1-Pet1* neurons. In previous work, we sorted fluorescently labelled mr*En1-Pet1* neurons followed by single-cell transcriptomic analyses^7^, which revealed that mr*En1-Pet1* neurons not only express the expected *Tph2* gene, which encodes the rate limiting enzyme in 5-HT synthesis, but also express the *Vglut3* gene, which encodes the vesicular glutamate transporter type 3 for packaging glutamate into synaptic vesicles. This finding suggests that mr*En1-Pet1* neurons may transmit using glutamate as well as 5-HT; indeed, for other 5-HTergic neurons such as in the dorsal raphe, VGLUT3 has been demonstrated to be sufficient for glutamate neurotransmission^28,29^. In GFP-labelled mr*En1-Pet1* neurons, we confirmed presence of *Vglut3* transcripts using single molecule fluorescent *in situ* hybridization (RNAscope) coupled with GFP immunodetection (Fig. 1d-e), and, furthermore, detected VGLUT3 protein along with 5-HT immunoreactivity (Fig. 1f). We then mapped the *En1-Pet1* efferent boutons (en passant and terminal) illuminated via synaptophysin-GFP (Syp-GFP) in triple transgenic *En1-cre; Pet1-Flpe; RC-FPSit* mice (aka *En1*^*cre/+*^; *Pet1-Flpe*^*Tg/0*^; *Gt(ROSA)26Sor*_*tm10(CAG-Syp/EGFP1,-tdTomato)Dym/+*_) (Fig. 1b,g), finding dense innervation of the SCN, as expected from our previous work^24^ (Fig. 1h), but now applying super-resolution imaging, we found that the mr*En1-Pet1* boutons innervating the SCN harbour 5-HT but not VGLUT3. And while levels of 5-HT and mRNA for *Tph2* in general have been reported to vary over the sleep/wake cycle^30,31^, this VGLUT3 configuration we observed in tissue harvested at Zeitgeber (ZT) 2 (0900 hrs; highest sleep pressure) was indistinguishable from tissue harvested at ZT14 (2100 hrs; lowest sleep pressure) suggesting stability across the sleep/wake cycle (Fig. 1i-j).

**Fig. 1.**
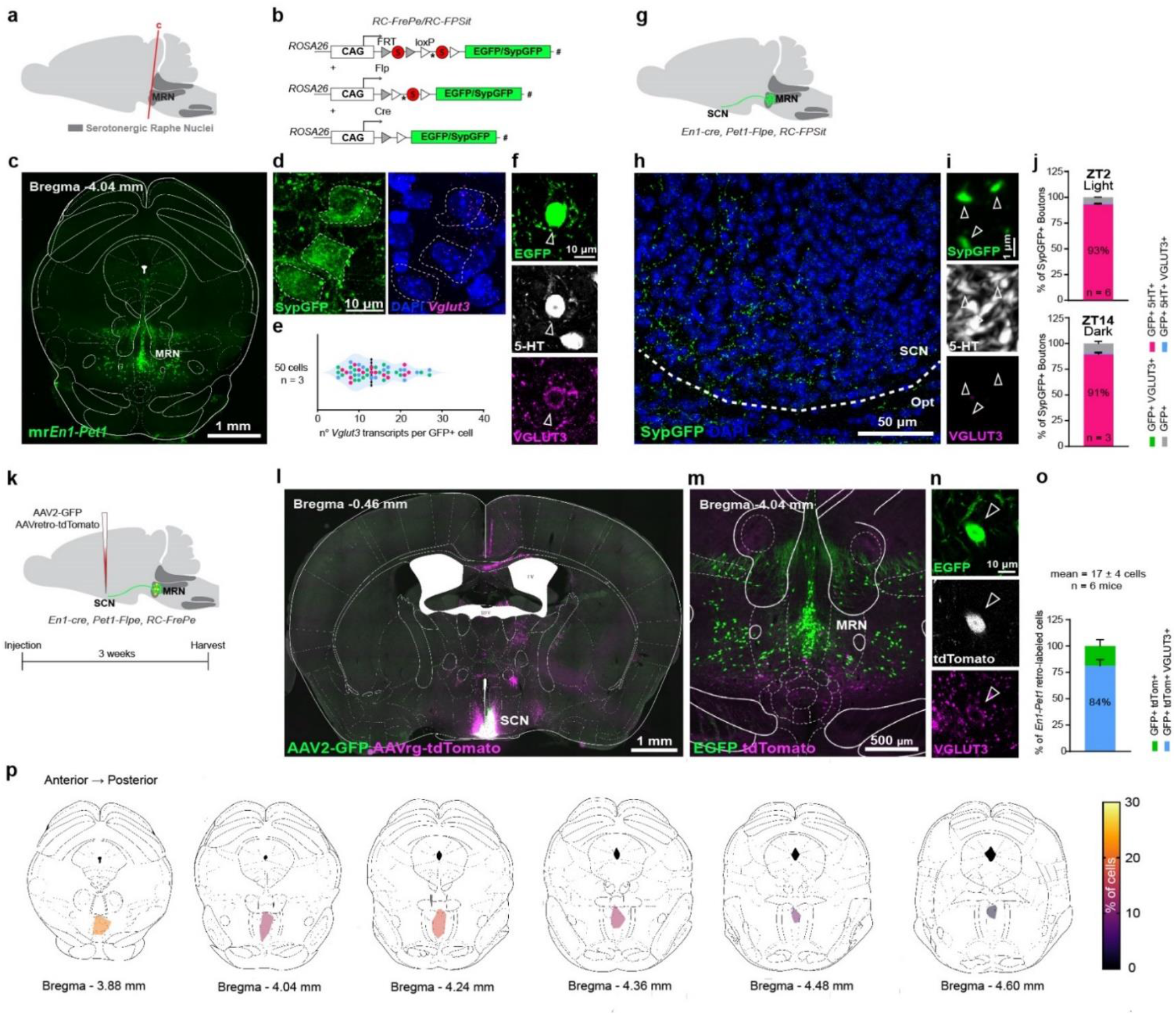
mr*En1-Pet1* soma harbour serotonin and VGLUT3 yet only serotonin is detected in the synaptic boutons innervating the SCN. **a**, Sagittal schematic representing the serotonergic raphe nuclei and location of the MRN. **b**, Intersectional genetic strategy and Cre- and Flp-dependent expression of EGFP (from *RC-FrePe* allele) and SypGFP (from *RC-FPSit* allele). * = mCherry cDNA in *RC-FrePe*. # = tdTomato cDNA in *RC-FPSit*. **c**, mr*En1-Pet1* neurons in the rostral MRN of *En1-cre;Pet1-Flp;RC-FrePe* mice. **d**, Dual IF-ISH showing *Vglut3* transcripts in mr*En1-Pet1* neurons (SypGFP^+^in *En1-cre;Pet1-Flp;RC-FPSit* mice). **e**, *Vglut3* transcripts per cell. Colors=different mice. Dotted line=mean. **f**, Triple IF showing costaining of 5-HT and VGLUT3 in GFP^+^ mr*En1-Pet1* neurons (in *En1-cre;Pet1-Flp;RC-FrePe* mice). **g**, Schematic showing projections from mr*En1-Pet1* neurons to the SCN. **h**, mr*En1-Pet1* neuron presynaptic boutons (SypGFP-labelled) innervating the SCN. Opt = optic nerve. **i**, Superresolution micrographs showing colocalization of SypGFP and 5-HT, but not with VGLUT3, in the SCN. **j**, Quantification of SypGFP^+^ boutons positive for 5-HT and/orVGLUT3 in the SCN assayed at two opposite time points of the circadian cycle. ZT = Zeitgeber. **k**, Retrograde tracing from the SCN. **l**, Representative image of the injection site. **m**, Representative image of retrogradely labelled cells in the MRN. **n**, SCN-projecting mr*En1-Pet1* soma show detectable VGLUT3. **o**, Quantification of the neurochemical phenotype of SCN-projecting mr*En1-Pet1* neurons. **p**, Heatmap showing the anatomical position of SCN-projecting mr*En1-Pet1* soma. All data expressed as mean ± s.e.m.

To corroborate the finding that mr*En1-Pet1* neurons lack VGLUT3 at their SCN terminals despite showing expression at the soma, we performed AAV-mediated retrograde tracing from the SCN, allowing direct comparison of soma and bouton neurochemical phenotype. We therefore infused a mixture of AAV-CAG-GFP (to mark the injection site) and AAVretro-CAG-tdTomato (which preferentially transduces projecting neurons^32^) in the SCN of triple transgenic *En1-cre; Pet1-Flp; RC-FrePe* mice (aka *En1*^*cre/+*^; *Pet1-Flpe*^*Tg/0*^; *Gt(ROSA)26Sor*_*tm8(CAG-mCherry,-EGFP)Dym/+*_) in which mr*En1-Pet1* neurons express EGFP (Fig. 1l, Extended Data Fig.1). We found that the mr*En1-Pet1* neurons that project to the SCN (mr*En1-Pet1*^*→SCN*^) – in this approach highlighted by the simultaneous presence of intersectionally-encoded EGFP and viral tdTomato – are a small subset within the broader mr*En1-Pet1* domain (Fig. 1m-n) and are preferentially located in the rostral MRN (Fig. 1p). ∽Eighty percent of the mr*En1-Pet1*^*→SCN*^ soma showed detectable VGLUT3 (Fig. 1o). We confirmed this finding using the retrograde dye Fluorogold infused in the SCN of *En1-cre, Pet1-Flpe, RC-FrePe* mice (Extended Data Fig. 2), ruling out synaptic bias by AAVretro. In both tracing experiments, no dorsal raphe *En1-Pet1* neuron projection to the SCN was observed, as expected.

Because serotonergic axons typically collateralize to multiple brain regions^33–35^, we explored the innervation profile specifically of the mr*En1-Pet1*^*→SCN*^ neuron subgroup. We infused a mixture of AAV-CAG-tdTomato and AAVretro-CAG-tdTomato in the SCN of *En1-cre; Pet1-Flpe; RC-FPSit* mice (Extended Data Fig. 3a-b). We found collaterals of mr*En1-Pet1*^*→SCN*^ neurons (double labeled with intersectional Syp-GFP and viral tdTomato) in the Paraventricular Nucleus of the Thalamus (PVT) and in the Septofimbrial Nucleus (Sfi) (Extended Data Fig. 3c). Notably, VGLUT3 was detectable in the collaterals to the Sfi but not PVT (Extended Data Fig. 3c). Next, we infused AAVretro-CAG-tdTomato in the PVT and separately Sfi of triple transgenic *En1-cre; Pet1-Flpe; RC-FPSit* mice and observed double-labelled SypGFP, tdTomato boutons in the SCN (Extended Data Fig. 3d-g), confirming this pattern of collaterals. These data indicate that dual transmitter mr*En1-Pet1*^*→SCN*^ neurons segregate 5-HT versus glutamate (proxy VGLUT3) to distinct collaterals innervating different brain regions, suggesting differential deployment of neurotransmitters in a branch- and region-specific manner.

### Shift from equinox to short/long photoperiod induces branch-specific reorganization of neurotransmitter deployment

Because changes in photoperiod duration and timing can induce switching of neurotransmitter usage by neurons in the SCN^6^, we queried whether neurotransmitter deployment by mr*En1-Pet1*^*→SCN*^ neurons might change upon shifting away from equinox to long or short photoperiod conditions. We exposed *En1-cre, Pet1-Flpe, RC-FPSit* mice, having previously been reared in equinox photoperiod conditions, to either long (19h light/5h dark) or short (5h light/19h dark) photoperiod for 2 weeks during adulthood^6,14,36^ (Fig. 2a) before harvesting brain tissue 2 h post light onset (light onset is fixed across conditions) for in situ phenotyping of mr*En1Pet1* boutons. We found that the shift to long photoperiod exposure increased the percentage of triple positive (SypGFP^+^–5-HT^+^–VGLUT3^+^) mr*En1-Pet1* boutons in the SCN up to 30% without changing the overall number of GFP^+^ boutons; and more strikingly, the shift to short photoperiod exposure increased the fraction of triple positive boutons up to 65% (Fig. 2b,d), again with the number of GFP^+^ boutons per field of view being unchanged across photoperiod conditions (Fig. 2c, f). When we exposed a separate cohort of *En1-cre, Pet1-Flpe, RC-FPSit* mice to short photoperiod for 2 weeks (Fig 2e), followed by standard equinox conditions for 2 weeks, we found that the percentage of triple positive mr*En1-Pet1* boutons in the SCN returned to previous equinox levels (<1%; Fig. 2g), suggesting reversibility of this VGLUT3 deployment.

**Fig. 2.**
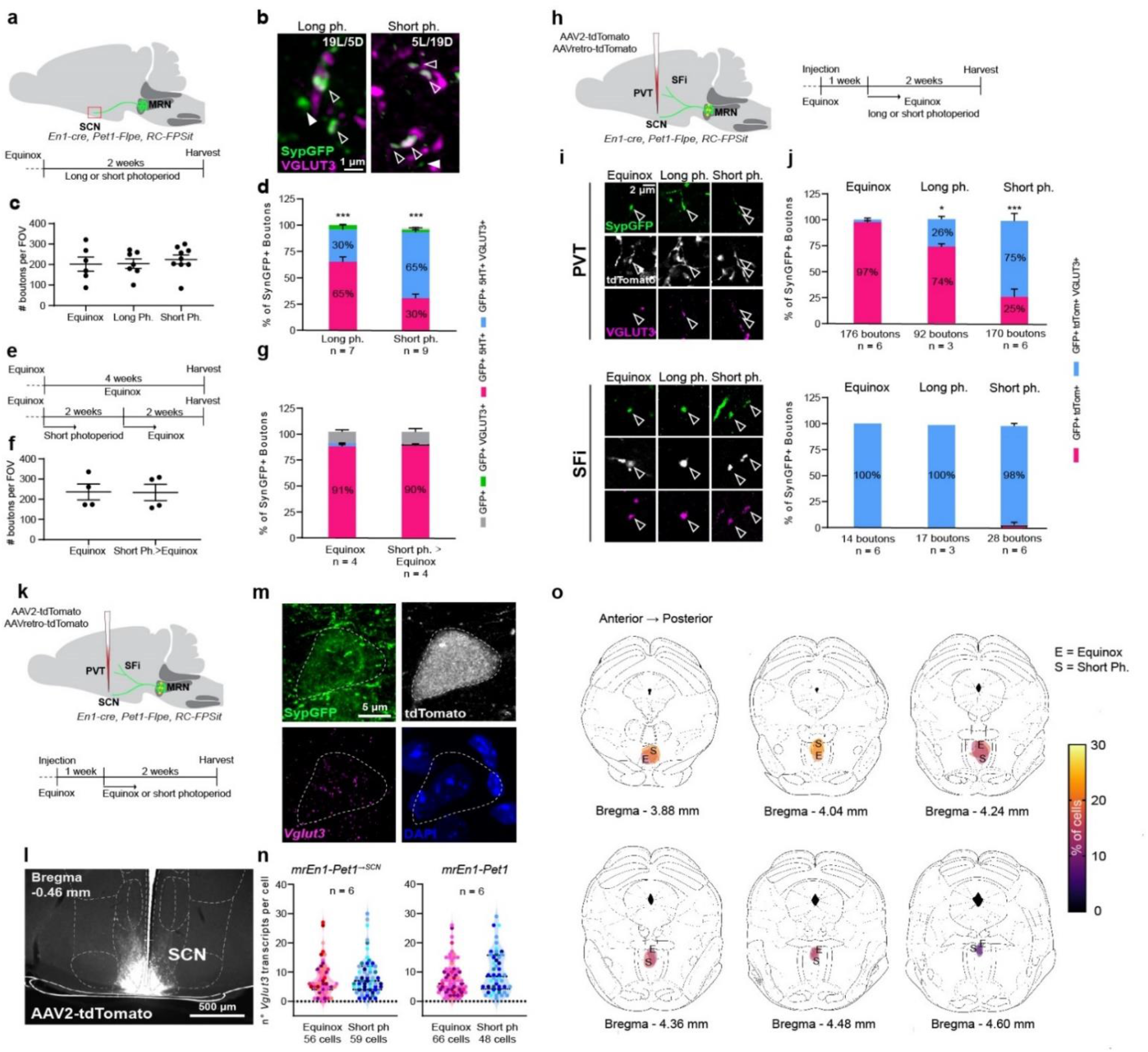
mr*En1-Pet1* neurotransmitter deployment reorganizes reversibly in a branch-specific manner in response to changes in photoperiod. **a, S**chematic of the experiment. SCN-innervating boutons from mr*En1-Pet1* neurons were analysed following 2 weeks of long/short photoperiod. **b**, Representative images showing VGLUT3 localization in mr*En1-Pet1* boutons in the SCN following photoperiod changes. **c**, Number of SypGFP^+^ boutons per field of view (FOV). **d**, Quantification of SypGFP-5-HT-VGLUT3 overlap in the SCN of *En1-cre;Pet1-Flpe;RC-FPSit* mice following shift to long versus short photoperiod (one-way ANOVA, *F* (2, 19) = 66.58, p < 0.0001. Equinox vs. Long ph. Tukey’s multiple comparison adjusted p = 0.0002; Equinox vs. Short ph. adjusted p < 0.0001; Long ph. vs. Short ph. adjusted p < 0.0001). **e**, Timeline to study reversibility. **f**, Number of SypGFP^+^ boutons per FOV. **g**, Quantification of SypGFP-5-HT-VGLUT3 overlap in the SCN of *En1-cre;Pet1-Flpe;RC-FPSit* mice in equinox and short ph.>equinox mice (two-tailed *t* test, t=1.644, df=6, p=0.15) **h**, Schematic of the experiment. Collaterals of mr*En1-Pet1*^*→SCN*^ neurons were analysed following 2 weeks of long/short photoperiod. **i**, Representative images showing VGLUT3 localization in mr*En1-Pet1* neuron collaterals innervating the PVT and SFi across photoperiod conditions. **j**, Quantification of SypGFP-5-HT-VGLUT3 overlap within PVT and SFi boutons in equinox, long and short photoperiod exposed mice (PVT: one-way ANOVA, *F* (2, 12) 6.720, p=0.011. Equinox vs. Long ph. Tukey’s multiple comparison adjusted p=0.0416; Equinox vs. Short ph. adjusted p<0.0001; Long ph. vs. Short ph. adjusted p=0.0004. SFi: one-way ANOVA, *F* (2,10) 0.5385, p=0.5997). **k**, Schematic of the experiment to study gene expression in mr*En1-Pet1*^*→SCN*^ neurons. **l**, Representative image of the injection site. **m**, Representative images showing individual *Vglut3* mRNA as puncta in mr*En1-Pet1*^*→SCN*^ soma. **n**, Quantification of *Vglut3* transcripts per cell in mr*En1-Pet1*^*→SCN*^ and mr*En1-Pet1* neurons in equinox-vs. short photoperiod-exposed mice. (mr*En1-Pet1*^*→SCN*^: two-tailed *t* test, t=0.8960, df=10, p=0.3910; mr*En1-Pet1*: two-tailed *t* test, t=0.1472, df=10, p=0.8859). **o**, Heatmap showing the anatomical position of SCN-projecting mr*En1-Pet1* neurons in equinox (E) vs. short photoperiod (S). All data expressed as mean ± s.e.m.

To study whether VGLUT3 deployment in PVT- and Sfi-innervating collaterals varies across photoperiods, we infused a mixture of AAV-CAG-tdTomato and AAVretro-CAG-tdTomato in the SCN of triple transgenic *En1-cre, Pet1-Flpe, RC-FPSit* mice and exposed them to either equinox, long, or short photoperiod for 2 weeks (Fig. 2h; Extended data Fig.4). PVT collaterals of mr*En1-Pet1*^*→SCN*^ neurons were devoid of VGLUT3 in equinox conditions; long photoperiod exposure increased the percentage of triple positive mr*En1-Pet1* boutons (SypGFP^+^, tdTomato^+^, VGLUT3^+^) to 25%, while the increase was ∽75% following short photoperiod exposure (Fig. 2i-j), mirroring the neurotransmitter re-organization observed in mr*En1-Pet1* boutons in the SCN. Sfi-innervating collaterals remained VGLUT3^+^, 5-HT^+^ double-positive across the different photoperiod conditions (Fig. 2i-j).

A possible mechanism by which mr*En1-Pet1*^*→SCN*^ neurons could augment VGLUT3 synaptic localization is through increased *Vglut3* gene transcription. To explore this possibility, we infused a mixture of AAV-CAG-tdTomato and AAVretro-CAG-tdTomato in the SCN of *En1-cre, Pet1-Flpe, RC-FPSit* mice, exposed them to either equinox or short photoperiod (Fig. 2k-l), and performed in situ detection of *Vglut3* transcripts concomitant with immunodetection of Syp-GFP (sufficient expression to illuminate soma) and viral tdTomato to distinguish the mr*En1-Pet1*^*→SCN*^ neurons. Transduced neurons showed comparable anatomical distribution between conditions (Fig. 2m, o) and analyses showed similar number of *Vglut3* transcripts per cell in mr*En1-Pet1*^*→SCN*^ neurons or other neighbouring mr*En1-Pet1* neurons (SypGFP^+^, tdTomato^-^, not projecting to the SCN), in response to short photoperiod exposure (Fig. 2n). This suggests that post-transcriptional (e.g. mRNA trafficking), translational, or post-translational (e.g. local translation, protein trafficking) mechanisms might underlie this reversible, photoperiod-induced neurotransmitter reorganization undertaken by mr*En1-Pet1*^*→SCN*^ neurons.

### Conditional loss of *Vglut3* in SCN-projecting neurons impairs adaptation to change in daylength

We hypothesized that this dynamic system of axon branch-specific neurotransmitter deployment by mr*En1-Pet1*^*→SCN*^ neurons is a cellular mechanism underlying organismal adaptation to change in photoperiod. Disrupting such deployment thus might cause circadian and sleep dysfunctions in response to changes in daylength, without affecting behaviour in equinox conditions. To test this hypothesis, we genetically disabled deployment of VGLUT3 in mr*En1-Pet1*^*→SCN*^ neurons and studied adaptation to different photoperiods as reflected in voluntary wheel running activity, the daily onset of which should rapidly synchronize to the new onset of darkness (Fig. 3a, b). Specifically, we infused a mixture of AAV2-CAG-GFP and either AAVretro-Syn-cre (experimental group) or AAVretro-CAG-tdTomato (control group) in the SCN of *Vglut3*^*flox/flox*^ mice^52^ to delete the floxed *Vglut3* gene in mr*En1-Pet1*^*→SCN*^ neurons. We found that all Cre-transduced (immunopositive) mr*Pet1* neurons (TPH2^+^) lacked VGLUT3 immunoreactivity (Fig. 3c, d), suggesting efficient deletion of the *Vglut3* gene by virally encoded Cre protein. By contrast, control animals transduced by AAVretro-tdTomato (instead of AAVretro-Syn-cre) showed, as expected, VGLUT3^+^ immunoreactivity in ∽half of the double positive TPH2^+^, tdTomato^+^ mr*Pet1* neurons (Fig. 3c, d). To explain the latter, it is important to note that this employed approach using current tools to maximize cell-selectivity for deletion of *Vglut3* nonetheless does so in all SCN-projecting neurons, thus not only mr*En1-Pet1*^*→SCN*^ neurons but also a serotonergic, non-glutamatergic (*Vglut3*^*-*^) subset of the MRN neuron group called *r2Hoxa2-Pet1*^7,37^. Across these two MRN neuron groups that project to the SCN collectively (mr*En1-Pet1*^*→SCN*^ and *r2Hoxa2Pet1*^*→SCN*^ neurons), only ∽50% of the soma express *Vglut3* transcript and VGLUT3 protein, i.e. the mr*En1-Pet1*^*→SCN*^ neuron subtype; *r2Hoxa2-Pet1*^*→SCN*^ neurons do not^37^ (corresponding to the VGLUT3-negative subset in AAVretro-tdTomato injected animals; Fig. 3d). We surmise that no other VGLUT3^+^ neuron population projects to the SCN because the VGLUT3^+^ boutons in the SCN always colocalize with 5-HT (across photoperiod conditions) and thus are from *Pet1* neurons. Collectively, these data suggest that the employed genetic approach results in functional loss of VGLUT3 selectively in mr*En1-Pet1*^*→SCN*^ neurons.

**Fig. 3.**
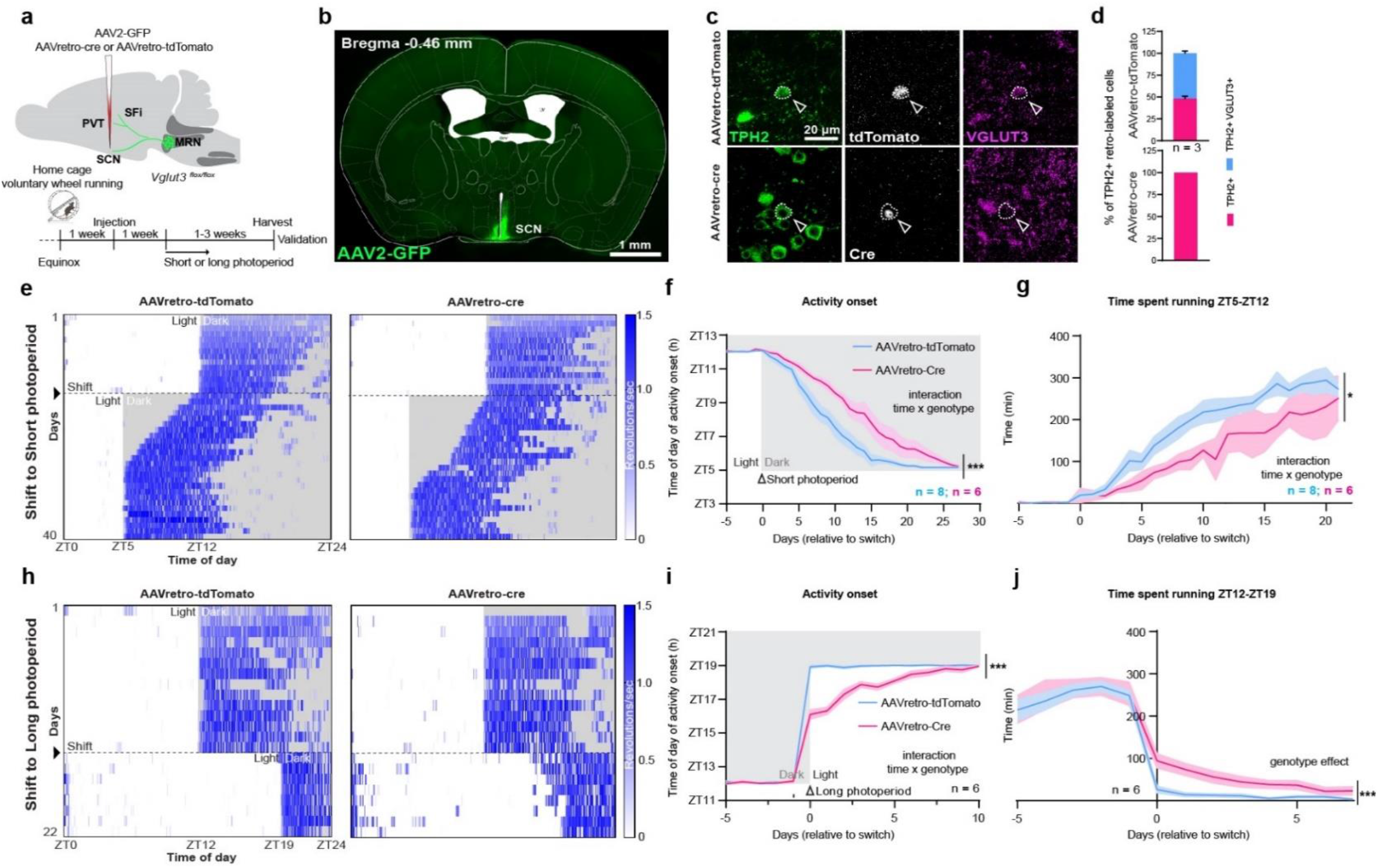
Genetic blockade of VGLUT3 deployment delays behavioural adaptation to changes in photoperiod. **a, S**chematic of the experimental approach and timeline. **b**, Representative image of the injection site. **c**, Efficient Cre-mediated knock-out of *Vglut3* in SCN-projecting serotonin neurons. **d**, Quantification of *Vglut3* knockout efficiency in SCN-projecting serotonin neurons. **e**, Representative heatmaps of control and cKO mice daily running activity (expressed as number of wheel revolutions per second) during habituation to short photoperiod. Black arrow and dashed black line indicate switch from equinox to short photoperiod. **f**, Activity-onset curve showing the time of day at which mice initiate running activity across days relative to short photoperiod switch (two-way ANOVA, interaction genotype x time *F* (32, 328) = 2.943, p<0.0001, followed by Sidak’s multiple comparison test). **g**, Time spent running between ZT5 and ZT12 (time window affected by light/dark manipulation) across days relative to short photoperiod switch (two-way ANOVA, interaction genotype x time *F* (26, 270) = 1.568, p=0.0425, followed by Sidak’s multiple comparison test. **h**, Representative heatmaps of control and cKO mice daily running activity during habituation to long photoperiod. Black arrow and dashed black line indicate switch from equinox to long photoperiod. **i**, Activity onset curve showing the time of day at which mice initiate running activity across days relative to long photoperiod switch (two-way ANOVA, interaction genotype x time *F* (12, 52) = 10.94, p<0.0001, followed by Sidak’s multiple comparison test. **j**, Time spent running between Z12 and across days relative to long photoperiod switch (two-way ANOVA, genotype effect *F* (1, 130) = 15.85, p = 0.0001, followed by Sidak’s multiple comparison test. Short photoperiod: n = 8 (control), 4 (cKO). Long photoperiod: n = 6 (control), 6 (cKO). All data expressed as mean ± s.e.m. Curve graph: solid line, mean; shaded area, s.e.m.

Next, we checked whether *Vglut3* deletion in SCN-projecting *Pet1* neurons would cause circadian disturbances in mice continuously exposed to equinox conditions. AAVretro-Cre-injected *Vglut3*^*flox/flox*^ (conditional knockout, cKO) mice showed no gross circadian abnormalities and were indistinguishable from control AAVretro-tdTomato-injected *Vglut3*^*flox/flox*^ mice (Extended data Fig. 5a-d). Next, we exposed a separate cohort of cKO and control mice to short photoperiod and monitored their habituation. Control mice advanced their activity onset (time of day at which they started running) over the course of the experiment, taking ∽15 days to phase synchronize to the new dark onset (Fig. 3e, f). By contrast, cKO mice showed significantly impaired phase synchronization, as illustrated by a much-delayed onset of activity across days relative to photoperiod switch (Fig. 3e, f), and less time overall spent running between ZT5 and ZT12 (daily time window affected by the light/dark shift; Fig. 3g) as compared to control mice. Overall, it took the cKO mice ∽25 days to fully synchronize their activity to the new dark onset. We next exposed a novel cohort of cKO and control mice to the long photoperiod. Control mice synchronized and restricted their locomotor activity to the new dark phase rapidly (light is a potent, aversive stimulus and bright environments pose risk for predation; Fig. 3h, i). By contrast, cKO mice showed bouts of activity during the light phase (Fig. 3i) and significantly greater time spent running between ZT12 and ZT19 (the daily time window affected by this photoperiod shift; Fig. 3j) that gradually reached control levels, suggestive of delayed adaptation. Further, we found significant correlation between the number of retrogradely Cre-transduced cells in the MRN, thus the number of mr*En1-Pet1*^*→SCN*^ neurons in which the *Vglut3* gene has been deleted, and the duration (number of days) required to habituate to both short and long photoperiod (Extended data Fig. 6). These data are consistent with the idea that timely and robust behavioural adaptation to photoperiod change requires *Vglut3* expression, allowing for neurotransmitter re-organization in mr*En1-Pet1*^*→SCN*^ neurons.

We next tested whether such genetic blockade of VGLUT3 deployment could also affect sleep/wake patterns in mice exposed to photoperiod change. Therefore, we infused a mixture of AAV2-CAG-GFP and either AAVretro-Syn-cre or AAVretro-CAG-tdTomato into the SCN of *Vglut3*^*flox/flox*^ mice, implanted them with electroencephalography/electromyography (EEG/EMG) probes connected to a wireless headmount for chronic unrestrained sleep/wake monitoring, and exposed them to either short or long photoperiod (Fig. 4a). cKO mice showed levels of wake, N-REM sleep, and REM sleep in baseline, equinox conditions comparable to that exhibited by control mice (Extended data Fig. 7a). Upon exposure to short photoperiod, control mice advanced their peak wake time, progressively synchronizing it to the new dark onset (Fig. 4b), and increased the time spent awake and simultaneously decreased time spent asleep between ZT5 and ZT12 (Fig. 4 c,d). Strikingly, cKO mice showed a completely blunted response to this photoperiod shift. Their peak wake time remained synchronized to the previous dark onset, and they showed no increase in wakefulness nor decrease in sleep between ZT5 and ZT12 (Fig. 4 c, d; Extended Data Fig. 7 c, d). We next exposed a separate cohort of cKO and control mice to long photoperiod and monitored their sleep/wake. As with locomotor behaviour, control mice quickly confined wake time to the lights-off period (ZT19-ZT0; Fig. 4e), decreased the overall amount of time spent in wakefulness, and increased the amount of sleep between ZT12 and ZT19 – the time window previously dark under equinox conditions (Fig. 4f, g). Thus, in this shift to long photoperiod, cKO mice did show adaptive responses (avoidance being a prevailing response to a light stimulus), however, their synchronization was delayed significantly as compared to control mice (Fig. 4e-g; Extended Data Fig, 7e, f) and cKO mice never reached control levels for overall wakefulness and sleep duration between ZT12 and ZT19, indicating aberrant sleep/wake timing.

**Fig. 4.**
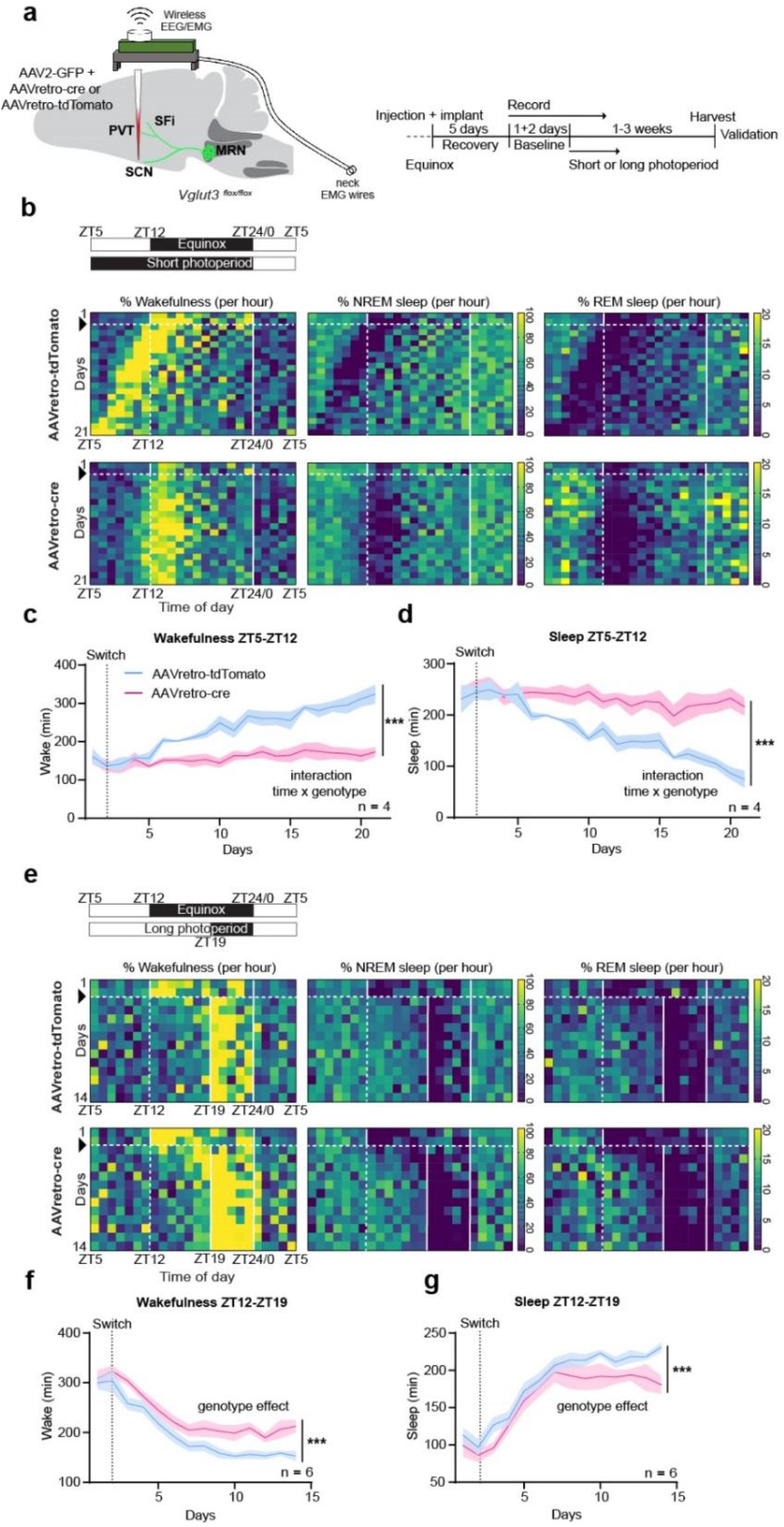
Genetic blockade of VGLUT3 deployment disrupts sleep/wake timing upon exposure to photoperiod change. **a, S**chematic of the experimental approach and timeline. **b**, Representative heatmaps depicting control (upper row) and cKO (lower row) daily wakefulness, NREM and REM sleep (expressed as % per hour) across days during habituation to short photoperiod (schedule schematized in upper left panel). **c**, Time spent awake between ZT5 and ZT12 (Mixed-effect model, interaction genotype x time, *F* (20, 114) = 9.757, p<0.0001, followed by Sidak’s multiple comparison test). **d**, Time spent asleep (NREM+REM) between ZT5 and ZT12 (Mixed-effect model, interaction genotype x time, *F* (20, 54) = 11.19, p<0.0001, followed by Sidak’s multiple comparison test). **e**, Representative heatmaps depicting control (upper row) and cKO (lower row) daily wakefulness, NREM and REM sleep across days during habituation to long photoperiod (schedule schematized in upper left panel). **f**, Time spent awake between ZT12 and ZT19 (two-way ANOVA, genotype effect, *F* (1, 139) = 56.78, p<0.0001, followed by Sidak’s multiple comparison test). **g**, Time spent asleep between ZT12 and ZT19 (two-way ANOVA, genotype effect, *F* (1, 145) = 29.22, p<0.0001, followed by Sidak’s multiple comparison test). Short photoperiod: n = 4 (control), 4 (cKO). Long photoperiod: n = 6 (control), 6 (cKO). Within the heatmaps, black arrow and horizontal dashed white line indicate photoperiod switch. Vertical dashed white line indicates time of day of previous dark onset. Vertical white solid lines delimit dark phase. All data expressed as mean ± s.e.m. Curve graph: solid line, mean; shaded area, s.e.m.

Thus, the result of this projection-specific genetic strategy to prevent neurotransmitter reorganization in mr*En1-Pet1*^*→SCN*^ neurons was disruption of locomotor and sleep/wake adaptation to changes in daylength.

### Monosynaptic inputs to mr*En1-Pet1* neurons

To begin delineating the input circuitry that engages mr*En1-Pet1* neurons to respond to photoperiod change, we performed intersectional, rabies-mediated trans-synaptic retrograde tracing^38^. We infused a 1:1 mixture of AAV8-Con-Fon-TVA-mCherry and AAV5-Con-Fon-oG in the MRN of double transgenic *En1-cre; Pet1-Flpe* mice (aka *En1*^*cre/+*^; *Pet1-Flpe*^*Tg/0*^), followed six weeks later by EnvA-pseudotyped, G-deleted Rabies Virus expressing GFP (RV-EnvA-DG-GFP). Brains were harvested one-week post-rabies injection, followed by immunofluorescence analyses and atlas registration for region specific RV^+^ (GFP^+^) cell counting (Fig. 5a, b). Validation analyses showed robust TVA-mCherry expression in Cre/Flp-double positive cells (Extended Data Fig. 8). However, similar to previous reports ^38^, we detected some mCherry in Cre-only cells, likely reflecting leaky expression from the antisense configuration of the Cre-recombined transgene; this prevented precise estimation of the starter cell population. No leaky expression was detected in Flp-only cells. Importantly, no trans-synaptic spread of the RV was detected in wild-type or single-transgenic animals^38^. Using this unbiased, brain-wide approach for mapping afferents, we identified four major monosynaptic inputs to mr*En1-Pet1* neurons: Preoptic Area (POA), Lateral Hypothalamus (LH), Lateral Habenula (LHb), and Ventral Tegmental Area (VTA) (Fig. 5c). Three of these regions (POA, LH, LHb) receive direct input from intrinsically photosensitive retinal ganglion cells and are known to influence sleep/wake behaviour as well as arousal^39–42^, thus representing possible circuit nodes capable of transmitting photoperiod information to mr*En1-Pet1* neurons.

**Fig. 5.**
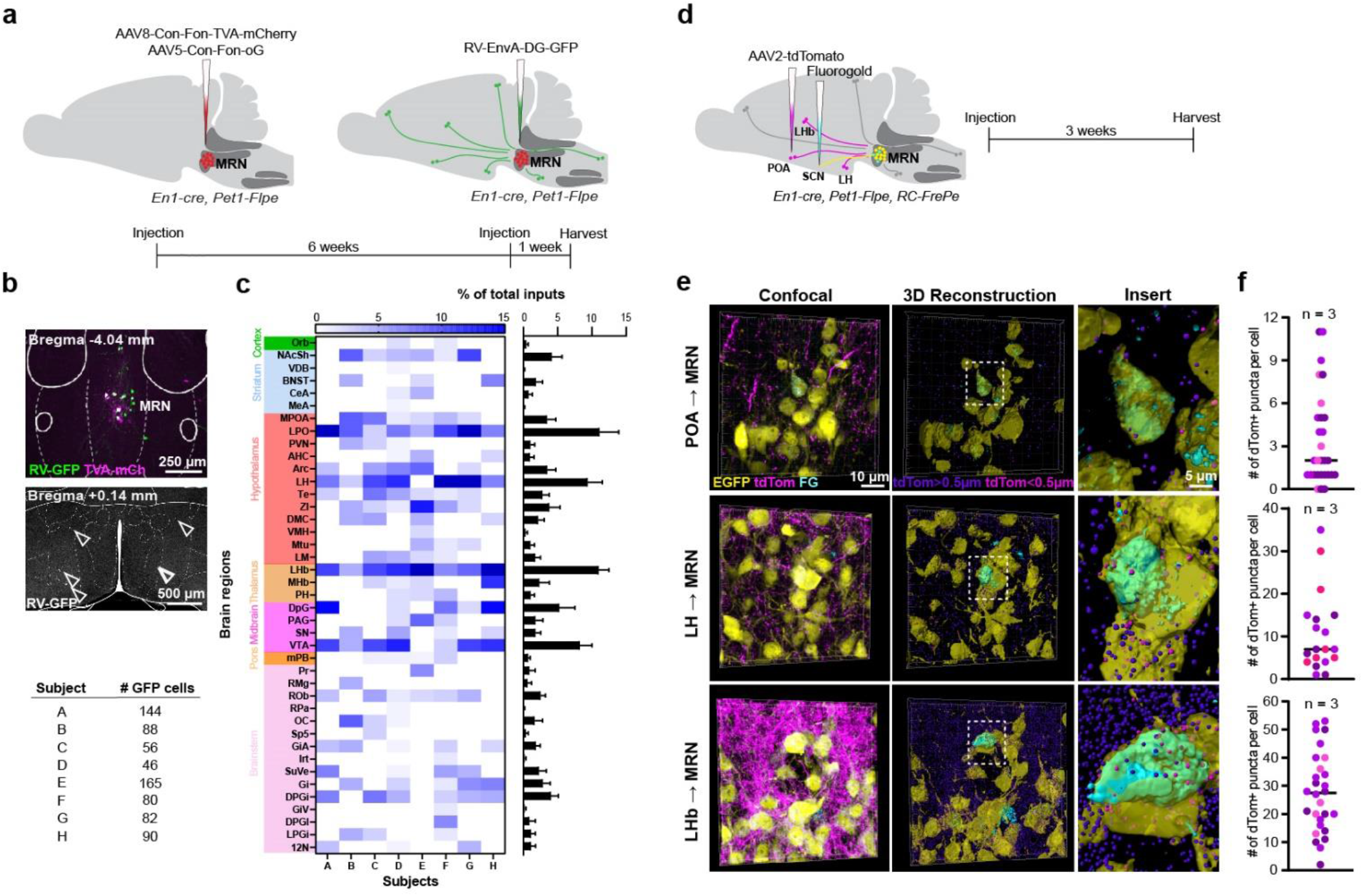
Brain-wide mapping of inputs to mr*En1-Pet1* neurons. **a**, Schematic of the experimental approach and timeline. **b**, Representative images of starter cells in the MRN (upper; scale bar 200μm) and retrogradely labelled upstream neurons in the POA (lower; scale bar 500μm). Total number of RV^+^ (GFP^+^) cells per mouse is reported in the bottom table. **c**, Heatmap showing atlas registration and quantification of input cells expressed as % of total inputs. Corresponding averaged data ± s.e.m. for each brain region is shown on the right. n = 8. **d, S**chematic of the experimental approach and timeline. **e**, Representative confocal images, and their 3D reconstructions, showing juxtaposition of boutons of input regions onto mr*En1-Pet1*^*→SCN*^ neurons. **f**, Quantification of the number of input boutons within 0.5μm of a mr*En1-Pet1*^*→SCN*^ neuron. Colors represent different mice. n = 3, for each input region.

Having delineated afferents to the mr*En1-Pet1* population as a whole, we next queried which provide input to the SCN-projecting subset of mr*En1-Pet1* neurons – i.e. to mr*En1-Pet1*^*→SCN*^ neurons. We therefore labelled mr*En1-Pet1*^*→SCN*^ neurons and their upstream inputs simultaneously and performed high-resolution imaging and 3D reconstructions to reveal those input boutons likely contacting mr*En1-Pet1*^*→SCN*^ soma. Specifically, we infused Fluorogold 4% in the SCN of *En1-cre, Pet1-Flpe, RC-FrePe* mice, highlighting mr*En1-Pet1*^*→SCN*^ neurons with GFP (intersectional) and Fluorogold, and infused the potential afferent source nuclei (POA, LH, or LHb, using separate cohorts) with AAV2-CAG-tdTomato (Fig. 5d). We segmented tdTomato-positive axons (coming from the input region) differentially based on their distance to mr*En1-Pet1*^*→SCN*^ neuron somata (GFP^+^ Fluorogold^+^) with a threshold of 0.5μm^43^ (Fig. 5e) and thus likely making synaptic contacts, which we observed from all three input regions (Fig. 5f), suggesting that the mr*En1-Pet1*^*→SCN*^ neuron group receives inputs from the POA, LH and LHb.

### POA→MRN circuit underlies adaptation to photoperiod shifts

Among the mapped inputs to mr*En1-Pet1*^*→SCN*^ neurons, we speculated the existence of at least one whose function is to decode and propagate photoperiod information to associated brain circuits, ultimately modulating adaptation. We reasoned that altering the activity of such a pathway might artificially mimic a change in daylength and consequently alter the percentage of 5-HT^+^ boutons in the SCN that are now also positive for VGLUT3^+^, much like that observed on shift from equinox to short/long photoperiod (Fig. 2b-d). We therefore devised an intersectional strategy for stable, non-invasive silencing of projection-defined neuronal populations *in vivo*. We infused AAVretro-cre in the MRN to deliver the *cre* transgene and thus Cre expression to input neurons, such as from the POA, LH, or Hb, and injected AAV-DIO-Flpo in either the POA, LH, or Hb of *RC-PFtox*^27^, *Ai65* trans-heterozygous mice (Fig. 6a; Extended Data Fig. 9). This compound intersectional configuration resulted in expression of the light chain of tetanus toxin (encoded by the *RC-PFtox* allele and expressed in a dual Cre/Flp-dependent fashion) and the reporter tdTomato (encoded by the *Ai65* allele, also expressed in a dual Cre/Flp-dependent fashion) selectively in either POA-, LH-, or Hb neurons (dual Cre^+^, Flpo^+^) projecting to the MRN, thus achieving stable and long term silencing of neuronal activity through tox-mediated inhibition of vesicular neurotransmission (Fig. 6a-c; Extended Data Fig. 9a-f). Notably, chronic silencing of POA→MRN neurons caused a significant increase of VGLUT3 and 5-HT colocalization within presynaptic boutons innervating the SCN as compared to controls (Fig. 6 c, d), whereas inactivating LH→MRN and Hb→MRN neuronal pathways showed no discernible effect on the percentage of VGLUT3 and 5-HT colocalization in the SCN (Extended Data Fig. 9a-f). These data suggest that POA, but not LH or Hb, neurons might convey to mr*En1-Pet1* neurons photoperiod information relevant to VGLUT3 deployment.

**Fig. 6.**
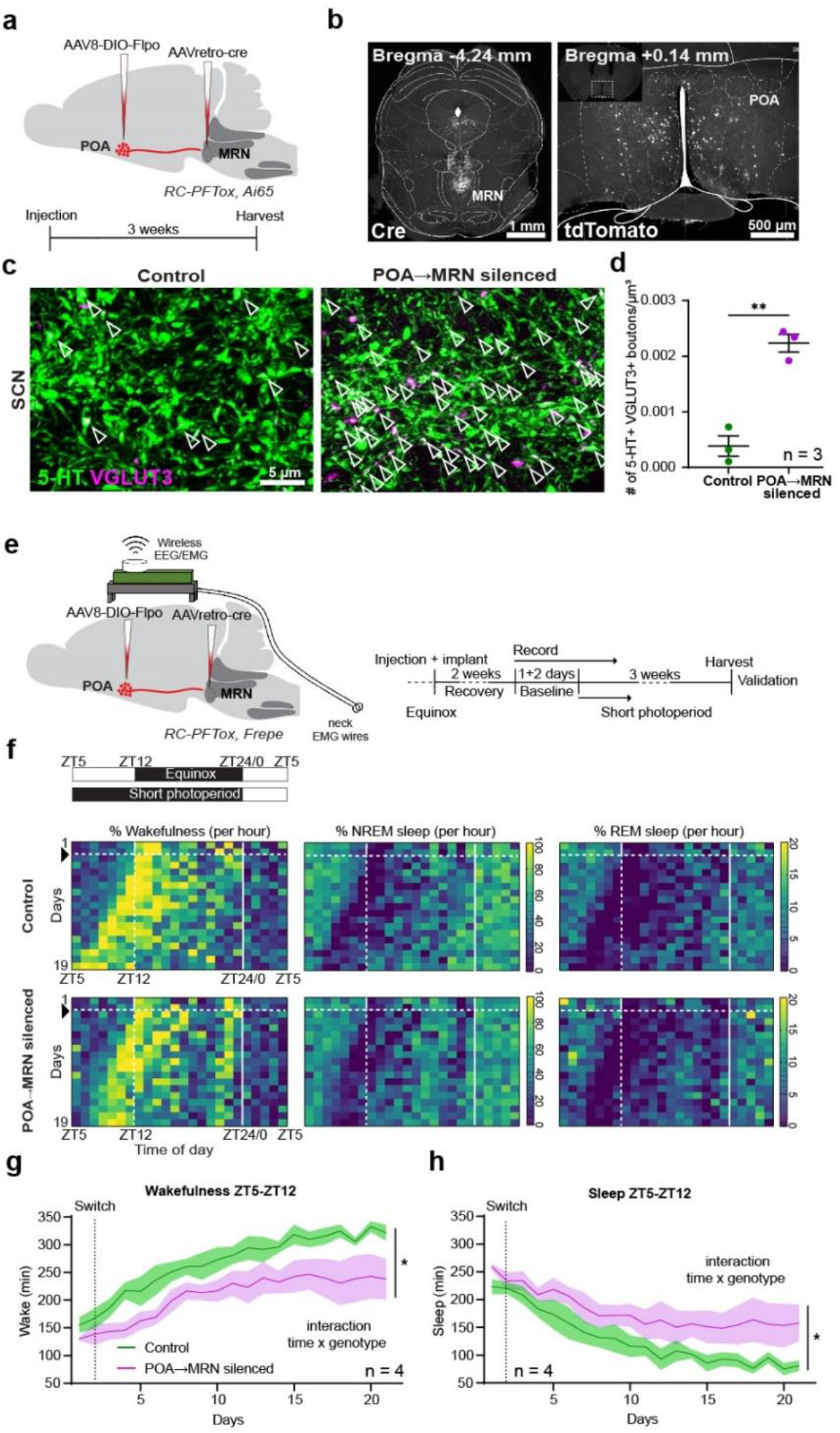
POA neurons projecting to the MRN convey photoperiod signals and modulate adaptation to changes in daylength. **a**, Schematic of the experimental approach and timeline. **b**, Representative image of the injection site in the MRN (AAVretro-cre; right) and in the POA (AAV-DIO-Flpo; left) where POA→MRN neurons are labelled with tdTomato (from dual Cre/Flpo-mediated recombination of the intersectional reporter Ai65). Scale bar, 1mm (left); 500μm (right). **c**, Superresolution images showing VGLUT3 localization within 5-HT axons in the SCN of mice in which POA→MRN have been silenced, as compared to control mice. Arrowheads indicate double positive boutons. **d**, Quantification of VGLUT3-5-HT colocalization in the SCN (two-tailed *t* test, t = 7.611, df = 4, p = 0.0016, n = 3). **e**, Schematic of the experimental approach and timeline. **f**, Representative heatmaps depicting control (upper row) and POA→MRN-silenced (lower row) mouse daily wakefulness, NREM and REM sleep across days during habituation to short photoperiod (schedule schematized in upper left panel). Black arrow and dashed white line indicate switch to short photoperiod **g**, Time spent awake between ZT5 and ZT12 (two-way ANOVA, interaction time x genotype, *F* (20, 60) = 2.027, p = 0.0186, followed by Sidak’s multiple comparison test). **h**, Time spent asleep (NREM+REM) between ZT5 and ZT12 (two-way ANOVA, interaction time x genotype, *F* (20, 60) = 1.978, p = 0.0317, followed by Sidak’s multiple comparison test). Short photoperiod: POA→MRN silenced). All data expressed as mean ± s.e.m. Curve graph: solid line, mean; shaded area, s.e.m.

To directly test the idea that POA neurons might propagate photoperiod information to mr*En1-Pet1* neurons, we examined whether silencing POA→MRN neurons would alter adaptation to changes in day length. We therefore monitored sleep/wake behaviour in POA→MRN-silenced mice during exposure to short photoperiod. We infused AAVretro-cre in the MRN and AAV-DIO-Flpo in the POA of *RC-PFTox; RC-FrePe* trans-heterozygous mice and implanted them with EEG/EMG probes connected to a wireless headmount. Notably, while no sleep/wake difference was observed in equinox conditions (Extended Data Fig. 10a), POA→MRN-silenced mice showed delayed adaptation to short photoperiod as compared to control mice (Fig. 6 f). Silenced mice also showed reduced time spent awake and increased time spent asleep between ZT5 and ZT12 (Fig. 6g, h; Extended Data Fig. 10b-c), suggestive of disrupted habituation. Taken together, these data suggest that POA neurons are uniquely positioned to deliver to the MRN information related to light exposure, and that this neural circuit plays a critical role in adaptation to changes in photoperiod.

## Discussion

The neural capacity to decode change in daylength and accordingly adjust activity, behaviour, and sleep/wake timing is vital to the survival and well-being of most species^1^. Misalignment can lead to unproductive, energetically wasteful, and even life-threatening animal behaviours such as mistimed foraging or hunting that is unsuccessful and a predation risk, and for humans clinically, can increase risk for and worsen outcome of many diseases, from affective^2,3^ to metabolic^12^. Our study reveals in mice a novel cellular and circuit mechanism in the brain for detecting photoperiod change and phase-shifting sleep-wake and activity correspondingly. We propose that in humans this network control point may be engaged in response to changes in light exposure that include not only seasonal daylength variations, but also trans meridian travel, shift work, and screen time.

Central to this phase-detection and behavioural-correction network is a previously unappreciated form of axon 5-HT-VGLUT3 branch-specific neurotransmitter deployment, here exhibited by a subset of brainstem mr*En1-Pet1* serotonergic neurons that collateralize to circadian and sleep timing brain centres, the SCN, PVT, and SFi. Few but compelling examples have shown that specialized neuronal cells can segregate neurotransmitters to different endings^44,45^, suggesting differential, target-specific modulation^46,47^. Here we provide evidence that segregation is malleable and reversible, being shaped by environmental cues, and importantly that this remodelling is physiologically relevant. When we deleted the *Vglut3* gene in SCN-projecting neurons, disabling genetically this VGLUT3 reorganization in mr*Pet1* neurons, adaptation to the new photoperiod was dramatically impaired: Sleep/wake patterns either failed to synchronize to the new photoperiod or were aberrant, and synchronization of voluntary exercise/activity to the new photoperiod was much delayed, even under light conditions which in the wild would be strongly aversive given predation risk. Post-transcriptional mechanisms likely underlie the differential deployment of VGLUT3, given that *Vglut3* transcript levels in mr*En1-Pet1*^→SCN^ neurons as well as in other mr*En1-Pet1* neurons were indistinguishable before and after photoperiod shift. Rearranging the localization of segregated neurotransmitters thus represents an essential, non-canonical form of plasticity that expands mechanisms employed by neurons to finely modulate circuits and ultimately behaviour depending upon state and condition.

Interestingly, a more severe desynchronization to short photoperiod was unmasked when monitoring sleep/wake synchronization as compared to wheel running activity. This suggests that the latter might boost synchronization, in keeping with previous reports^48^. Further, sleep/wake desynchronization was milder in response to the long as compared to short photoperiod shift, indicating that avoidance to light might be a prevailing stimulus that induces mice to seek shelter and sleep more. Mechanistically, photoperiod-dependent neurotransmitter reorganization and resulting glutamate increase synaptically through augmented VGLUT3 localization could represent a reinforcing signal that works synergistically with environmental light input to reset clock neurons in the SCN. It is noteworthy that both short and long photoperiod shifts induce an increase in VGLUT3 within mr*En1-Pet1* boutons, albeit to different extents. We speculate that the postsynaptic target neurons may be different for those mr*En1-Pet1* boutons that become VGLUT3^+^ in response to a short vs. long photoperiod, thus modulating the circuit differentially depending on daylength. Interestingly, some SCN neurons (those positive for Vasoactive Intestinal Polypeptide and Neuromedin S) display adaptive neurotransmitter switching in response to photoperiod change^6^. Investigating whether mr*En1-Pet1* neurons contact SCN “switching” neurons, perhaps selectively via boutons that undergo VGLUT3 reorganization, and whether they play a causal role in this switching is an important future direction.

At the circuit level, we identified the POA as an essential node^49,50^ upstream of mr*En1-Pet1* neurons responsible for decoding and propagating photoperiod information. Adult-induced silencing of POA→MRN neurons was sufficient to cause neurotransmitter reorganization in 5-HT boutons innervating the SCN, artificially mimicking a change in daylength duration in otherwise equinox conditions; however, sleep/wake was unaltered. These findings suggest that deployment of VGLUT3 by mr*En1-Pet1*^→SCN^ neurons is not sufficient *per se* to modify sleep/wake patterns; albeit it is necessary, as shown by the *Vglut3* cKO results. Only when exposed to short photoperiod, POA→MRN-silenced mice showed disrupted sleep/wake adaptation, suggesting necessity of an intact POA-MRN circuit. The POA appears to be uniquely positioned anatomically to modulate photoperiod adaptation as it receives innervation from both retinal ganglion cells^39^ and SCN neurons^10^, thus representing an integration site of both direct light input and central clock output. Further circuit mapping will be required to discern whether such integration happens in individual or separate but locally interconnected POA cells, and whether these cells are those that innervate the MRN.

In summary, we identified a novel POA→mr*En1-Pet1*→SCN/PVT/SFi circuit essential for behavioural adaptation to changes in atmospheric light duration. A critical component of this circuit is the MRN mr*En1-Pet1* neuron group, which shows a novel form of periodic, branch-specific neurotransmitter reorganization that tracks with photoperiod changes and is necessary for behavioural adaptation. By identifying relevant molecules, cells, and circuit nodes, this work has the potential to inform therapeutic interventions for disorders characterized by circadian and sleep timing disturbances.

## Data availability

All data is available upon request.

## Acknowledgments

The authors are grateful to members of the Dymecki laboratory for discussions related to this work and manuscript, to the Micron Microscopy Core at Harvard Medical School and the Harvard Center for Biological Imaging (HBI) at Harvard Faculty of Arts and Science for infrastructure and support, as well to the Weitz laboratory at Harvard Medical School for sharing with us their circadian cabinets for pilot work foundational to this study. The work of the authors is supported by US National Institutes of Health grant (R21MH127341) to G.M. and S.M.D., the Baszucki Brain Research Fund with the Milken Institute Center for Strategic Philanthropy (grant to G.M. and S.M.D.), and the Harvard Brain Science Initiative (HBI) Bipolar Disorder Seed Grant Program supported by the Dauten Family Foundation and Petri Deryng (grant to G.M., R.A.S., and S.M.D.).

## Author contributions

G.M. and S.M.D. conceived the study and designed experiments. G.M. performed experiments. G.M. and Y.C. performed dual immunofluorescence *in situ* hybridization experiments. R.A.S. helped establish the wheel-running assay in the lab and wrote custom script for visualizing and analysing the activity data.

G.M. and S.M.D. wrote the manuscript.

## Methods

### Animals

All experimental procedures were approved by the Harvard Institutional Animal Care and Use Committee. Mice were raised and housed on a 12 h light/dark cycle, unless otherwise specified, under constant temperature (22±1°C) and humidity (40-50%). For short, long, and reverse-to-equinox photoperiod studies, mice were transferred to a dedicated, neighbouring room with independent light/dark control and ZT0 always at 0700 h, and exposed to either 19 h light/5 h dark cycle (short photoperiod), 5 h light/19 h dark cycle (long photoperiod), or 12 h light/12h dark (equinox) for at least 2 weeks, according to previously published protocols^6,14,36^. Mice of both sexes (equal numbers) were 60-70 days old at the start of every experimental procedure. All transgenics were backcrossed to the C57BL/6J strain (#000664, Jax Labs) for at least 9 generations. *En1*^*tm2(cre)wrst/J*^ mice^51^, referred to as *En1-cre* (Jax Lab #007916) were provided by A. Joyner. *Slc17a8*^*flox/flox*^ mice^52^, referred to as *Vglut3*^*flox/flox*^, were provided by R. Seal. *Gt(ROSA)26Sor*^*tm65*.*1(CAG-tdTomato)Hze*^/^J^ mice, referred to as *RC-Ai65* or *R26*^*C- Ai65/+*,^ were obtained from Jax Lab, stock #021875. Previously generated in the Dymecki lab were: Tg(*Fev-flpe)*1Dym mice, referred to as *Pet1-Flpe*^23^; *Gt(ROSA)26Sor*^*tm8(CAG-mDherry,-EGFP)Dym*^ mice, referred to as *RC-FrePe*^25^ or *R26*^*C-FrePe/+*^ (Jax Lab # 029486); *Gt(ROSA)26Sor*^*tm10(CAG-Syp/EGFP*,-tdTomato)Dym*^ mice, referred to as *RC-FPSit*^26^ or *R26*^*C-FPSit/+*^ (Jax Lab #0030206); and *Gt(ROSA)26Sor*^*tm(CAG-GFPTox*)Dym/J*^ mice, referred to as *RC-PFtox*^27^ or *R26*^*C-PFtox/+*^. Experimental triple transgenic *En1*^*cre/+*^, *Pet1-Flpe*^*Tg/0*^, *R26*^*C-FrePe/+*^ mice or *En1*^*cre/+*^, *Pet1-Flpe*^*Tg/0*^, *R26*^*C-FPSit/+*^ mice were generated by crossing double *En1*^*cre/+*^, *Pet1-Flpe*^*Tg/0*^ studs to homozygous reporter *R26*^*C- FrePe/C-FrePe*^ *or R26*^*C-FPSit/C-FPSit*^ females. *Vglut3*^*flox/flox*^ mice were generated by crossing *Vglut3*^*foxlflox*^ studs to *Vglut3*^*flox/flox*^ females. Double transgenic mice *En1*^*cre/+*^, *Pet1-Flpe*^*Tg/0*^ mice were generated by crossing *En1*^*cre/+*^, *Pet1-Flpe*^*Tg/0*^ studs to C57BL/6J females (#000664, Jax Labs). Trans heterozygotes *R26*^*C-PFtox/Ai65*^ or *R26*^*C-PFtox/C-*^ _*FrePe*_ were generated by crossing either *R26*_*C-FrePe/C-FrePe*_ or *R26*_*C-Ai65/C-Ai65*_ studs to *R26*_*C-PFtox/C-PFtox*_ females.

### Tissue preparation, immunofluorescence, and *in situ* hybridization

Histological analyses were performed on tissue harvested 2 h post light onset (Zeitgeber (ZT) 2 = 0900 h); light onset (ZT 0) was fixed at 0700 h regardless of photoperiod length, which is varied by changing lights-off timing at either 1900 h (ZT 12) for equinox, 1200 h (ZT 5) for short photoperiod, or 0200 h (ZT 19) for long photoperiod), thus controlling for phase, except for the ZT 14 cohort of Fig. 1j). Mice were perfused transcardially with 0.1 M PBS, followed by 4% Paraformaldehyde (PFA). Brains were dissected out and post-fixed in 4% PFA overnight (20-24h) at 4°C. Tissue was either embedded in 2.5% agarose and vibratome sectioned (Leica VT1000S) or cryoprotected in 30% sucrose, embedded in tissue freezing medium (Tissue Tek, OCT), and then cryo-sectioned (Leica): Twenty μm cryosections were used for dual immunofluorescence *in situ* hybridization, 50 μm vibratome sections for immunofluorescence followed by wide-field or confocal imaging, 20 μm vibratome sections for immunofluorescence followed by superresolution imaging.

For immunofluorescence experiments, tissue sections were permeabilized for 1h at room temperature in 0.1 M PBS containing 0.5% Triton X-100 (Sigma Aldrich), followed by 1.5 h incubation in blocking solution of 0.1 M PBS containing 0.3% Triton X-100 and 5% Normal Donkey Serum (NDS, Jackson Immunoresearch). Sections were then transferred in blocking solution containing primary antibodies and incubated for 1-2 overnights at 4°C. After 3 washes in 0.1 M PBS containing 0.3% Triton X-100, sections were incubated in blocking solution containing secondary antibodies for 2h at room temperature. Finally, sections were washed 3 times in 0.1 M PBS containing 0.3% Triton X-100 solution, mounted onto 1.5 glass coverslips (Electron microscopy science) and attached to Superfrost slides (ThermoFisher) using Prolong Gold mountant (ThermoFisher) for widefield and confocal imaging or Prolong Glass mountant (ThermoFisher) for superresolution imaging. We used the following primary antibodies: anti-GFP (chicken polyclonal, 1:1000, Aves Labs GFP-1020); anti-5-HT (goat polyclonal, 1:500; Abcam #ab66047); anti-VGLUT3 (guinea pig polyclonal, 1:500, Synaptic Systems #135–204); anti-RFP (rat monoclonal, 1:1000, Chromotek #5f8 used to stain for virally-expressed tdTomato only in the dual immuno-*in situ* while for all other experiments endogenous tdTomato fluorescence was visualized); anti-Fluorogold (rabbit polyclonal, 1:3000, Millipore AB153-I); anti-Cre (rabbit polyclonal, 1:500, Biolegend #908001); anti-TPH2 (goat polyclonal, 1:1000, Abcam #ab121013); and species-matched donkey secondary antibodies (1:500) conjugated with Alexa Fluor 488, Alexa Fluor 568, Alexa Fluor 647 (Invitrogen, Molecular probes) or Cy5 (Jackson Immunoresearch, #AB_2340462).

Combined immunofluorescence single molecule *in situ* hybridization (RNAscope) was performed using a two-step protocol. RNAscope was performed according to manufacturer instructions (RNAscope, Advanced Cell Diagnostics) using 20 μm sections mounted onto SuperFrost Plus Slides (ThermoFisher) hybridized with a *Vglut3* probe. Afterwards, sections were immunostained for GFP, or for GFP and RFP, using standard protocols as described above. For combined immunofluorescence RNAscope on mr*En1-Pet1*^→SCN^ neurons, sections were screened beforehand for the presence of retrogradely labelled tdTomato^+^, SypGFP^+^ neurons in the MRN as assessed by direct fluorescence using a widefield microscope. Only sections that showed positive cells were mounted onto slides and processed.

### Wide-field, confocal, and superresolution imaging and 3D analysis

Wide-field microscopy was used to acquire tiled whole-section images (mr*En1-Pet1* neuron anatomical location, AAV injection sites, Rabies Virus cell counting) using a Nikon Ti inverted microscope equipped with a Lumencor SOLA LED light engine illumination, Hamamatsu ORCA-Flash4.0 V3 digital CMOS camera, and a Plan Apo Lambda 10x/0.45 Air Dic N1 objective. Confocal microscopy was used to perform imaging of soma neurochemical phenotype, dual immunofluorescence-*in situ* hybridization, as well as validation of rabies tracing experiments using a Zeiss Axio Observer Z1 equipped with LSM780, Quasar PMT and Gasp 32 channels spectral detectors with a Plan Apo 63X/1.4 Oil DIC III objective. Voxel size was set to 0.13×0.13×0.31μm, resolution to 1024×1024. Superresolution imaging was used to assess the neurochemical phenotype of mr*En1-Pet1* boutons using either a Zeiss Axio Observer Z1 equipped with LSM980 scan head and Airyscan2 detector and Plan Apo 63x/1.4 Oil DIC III objective, or a Zeiss Axio Observer Z1 equipped with LSM880 Scan Hear with Airyscan detector and Plan Apo 63x/1.4 Oil DIC III objective using Superresolution imaging mode. Voxel size was set to 0.04×0.04×0.22 μm; resolution to 3148×3148; number of *z* planes to 23.

3D colocalization of SypGFP boutons with 5-HT and VGLUT3, with VGLUT3 and tdTomato, and between 5-HT and VGLUT3, was performed semiautomatically on Airyscan images using Imaris (Bitplane, v9) as previously described^37^. Briefly, each channel was segmented using the Spot function and artifacts were manually removed. Distance based colocalization was performed using the MATLAB plug-in “Colocalize Spots” with a threshold of 0.5 μm (distance between objects centroids).

Anatomical registration and quantification of Rabies virus GFP^+^ cells were performed by overlaying widefield microscopy images with the corresponding Paxinos Atlas table using Photoshop 2021. Cells were then assigned to specific brain regions and manually counted.

3D segmentation and quantification of input boutons onto mr*En1-Pet1*^→SCN^ neurons was performed semiautomatically on confocal images using Imaris. GFP^+^ mr*En1-Pet1* neuron cell bodies and Fluorogold immunostainings were segmented using the Surface function, while tdTomato^+^ boutons were segmented using the Spot function. The MATLAB plug-in “Spot and Surface Distances” was used to segment and calculate the number of tdTomato^+^ boutons that were closely associated with GFP^+^, Fluorogold^+^ cell bodies with a distance -o-soma threshold of 0.5 μm.

### Stereotaxic injections

All surgeries were performed under sterile conditions. Mice were anesthetized with isoflurane (4% induction, 1.5% maintenance) and placed in a stereotaxic apparatus (Kopf) equipped with a heating pad. After removing fur and cleaning the scalp with ethanol and betadine, an incision was made using a sterile scalpel. The skull was levelled anteroposteriorly between Bregma and Lambda and mediolaterally between two points equidistant 1.5 mm from Bregma, such that all four points were at 0.0 ± 0.1 mm *z* depth. After making a point-size craniotomy with a dental drill, viruses were delivered using a thin glass pipette (50 μm at the tip) connected to a Nanoject III (Drummond) at 60nL/min (100-200 nL). For SCN injections specifically, 3 steps (2 steps for Fluorogold [Fluorochrome] injections) of 5 nL each (1 min apart) were performed at 46 nL/sec speed. When the injection was completed, the pipette was left in place for an additional 10 min, and then slowly withdrawn. The skin wound was then closed with absorbable sutures. Mice were monitored up to 5 days post-operation and administered Meloxicam 10 mg/kg (once a day for three days) and slow-release Buprenorphine 1 mg/kg (once, administered after surgery) to minimize pain. We used the following coordinates relative to Bregma: SCN -0.4 mm anteroposterior (AP), ±0.15 mm mediolateral (ML), -5.8 mm dorsoventral (DV) unilaterally (though achieving bilateral spread); MRN -4.2 mm AP, 0.0 mm ML, -4.5 mm DV unilaterally; POA +0.14 mm AP, ±0.5 mm ML, -5.5 mm DV bilaterally; LH -1.58 mm AP, ±1.25 mm ML, -5.0 mm DV bilaterally; Hb -1.34 AP, ± 0.35mm ML, -2.5mm DV bilaterally. To account for differences in brain size, Bregma-Lambda distance of each experimental mouse was measured and divided by 4.2 mm (∽ Bregma-Lambda distance according to Paxinos Atlas) to calculate a normalizing factor that was used to scale the anteroposterior coordinate.

Mice showing incorrect viral targeting, or partial coverage of the target area, were excluded from the study.

## Viral vectors

The following viral vectors were used: AAVretro-CAG-tdTomato (Boston Children’s Hospital (BCH) Viral Core), 1.2 × 10^12^ genomic copies (gc)/mL; AAV2-CAG-GFP (BCH Viral Core), 2 × 10^12^ gc/mL; AAV2-CAG-tdTomato (BCH Viral Core), 1.8 × 10^11^ gc/mL; AAVretro-Syn-Cre (Addgene #105553, a gift from James M. Wilson) 4 × 10^12^ gc/mL; AAV8-Con-Fon-TVA-mCherry (Addgene plasmid #131779, a gift from Karl Deisseroth) and AAV5-Con-Fon-oG (Addgene plasmid #131778, a gift from Karl Deisseroth) were custom packaged by the BCH Viral Core at titers 3.2 × 10^13^ gc/ml and 2.4 × 10^13^ gc/ml, respectively; EnvA pseudotyped G-Deleted Rabies Virus GFP (Salk Institute GT3 Viral Core), 2 × 10^8^ infection units/mL; AAV8-DIO-Flpo (Addgene #87306-AAV8, a gift from Li Zhang) 2.3 × 10^12^ gc/mL.

### Home cage running-wheel assay

For the duration of the running-wheel experiments, mice of either sex were single housed. Running wheels were custom built by Ofer Mazor and Pavel Gorelik from the Harvard Medical School Instrumentations Core Facility. Briefly, metal wheels were mounted on a plexiglass transparent support that fit standard mouse cages. An infrared sensor was integrated within the plexiglass support to detect the spokes of the wheel passing by (a single revolution was considered when the two spokes of the wheel passed by the sensor) and connected to a 24-port Raspberry Pi running a custom-made script for activity detection. Data was expressed as revolutions/second, visualized and analysed in 10 sec bins using custom R scripts. Sta) and Prism 9 (GraphPad). Mice were allowed to habituate to the running wheel for at least a week before collecting activity data.

### Chronic sleep/wake measurements

Following stereotaxic injection of AAVs, mice were implanted with 2-channel Electroencephalography (EEG)/1-channel Electromyography (EMG) headmounts (Pinnacle Technology, # 8201-SS). Four screws (front 0.10’’, #8209; back 0.12’’, #8212, Pinnacle Technology) were inserted into the skull to anchor the headmount and simultaneously record cortical EEG signals. Two EMG wires positioned in the neck muscle recorded EMG signals. The headmount was secured using dental cement. Mice recoveredw for 5 days, and then were connected to the wireless EEG/EMG recording device (Pinnacle Technology, # 8200-K9-SL) for 24 hours (habituation) before the start of the recording session. Each device (up to 4 simultaneously, to avoid Bluetooth interference as per manufacturer recommendation) was then paired with a Bluetooth receiver connected via USB to a computer running the Sirenia data acquisition software (Pinnacle Technology). Baseline sleep/wake was recorded continuously over the following 2 days, before switching to either short or long photoperiod. Recordings lasted up to 3 weeks.

Analysis of sleep/wake was performed using Sirenia Sleep Pro (Pinnacle technology). Digitized polygraphic data were analysed in 10 sec bins, their power spectra generated and then classified as Wake, NREM and REM using the Cluster Scoring function. Data was then manually curated to either confirm/correct sleep/wake states or to score unidentified epochs. Sirenia Sleep Pro was used to calculate the amount of Wake/NREM/REM in defined time periods. Data was visualized using GraphPad Prism 9.

### Statistical analysis

Statistical tests (two-sided) were performed using GraphPad Prism 9. No statistical methods were used to predetermine sample size. Mean ± s.e.m. was used to report statistics. Significance was defined as p < 0.05. Details of the tests used, group numerosity and multiple hypotheses corrections can be found in the corresponding figure legends and in Multiple Comparisons Tables.

