## Extended Data figures for "A brain circuit and neuronal mechanism for decoding and adapting to change in daylength"

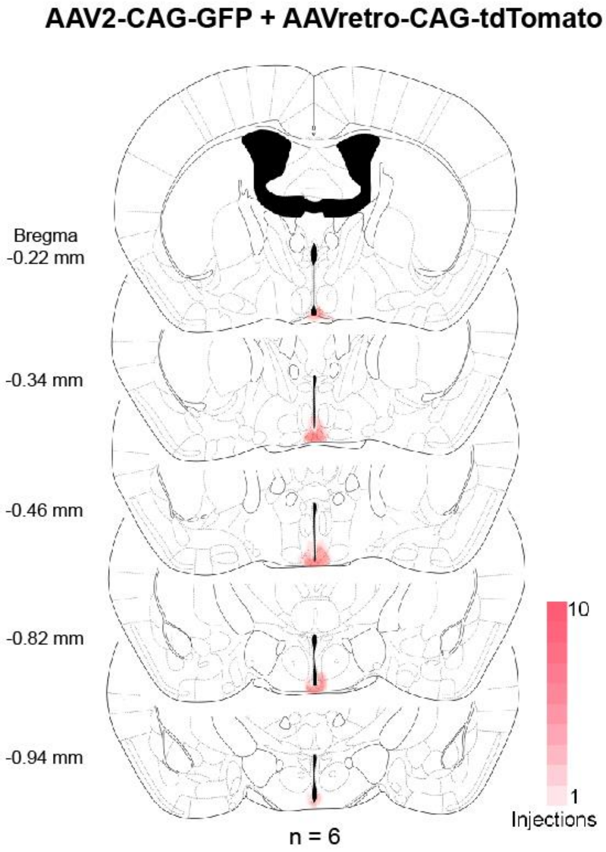


**Extended Data Fig. 1. Validation of retrograde viral circuit tracing from the SCN**. Schematic of color-coded viral targeting and spread across 5 coronal planes.

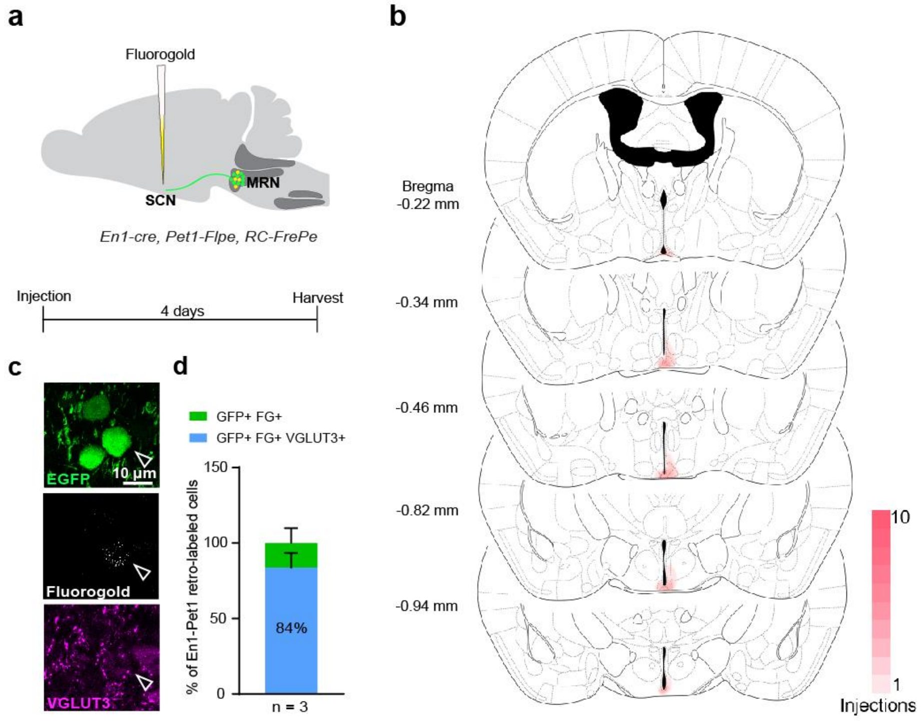


**Extended Data Fig. 2. Validation of retrograde, Fluorogold-mediated circuit tracing from the SCN. a,** Schematic of the experimental approach and timeline. **b,** Schematic of color-coded Fluorogold targeting and spread across mice. **c,** Representative image of retrogradely labelled cells in the MRN. SCN-projecting mr*En1-Pet1* neurons show detectable VGLUT3. **d,** Quantification of the neurochemical phenotype of SCN-projecting mr*En1-Pet1* neurons.

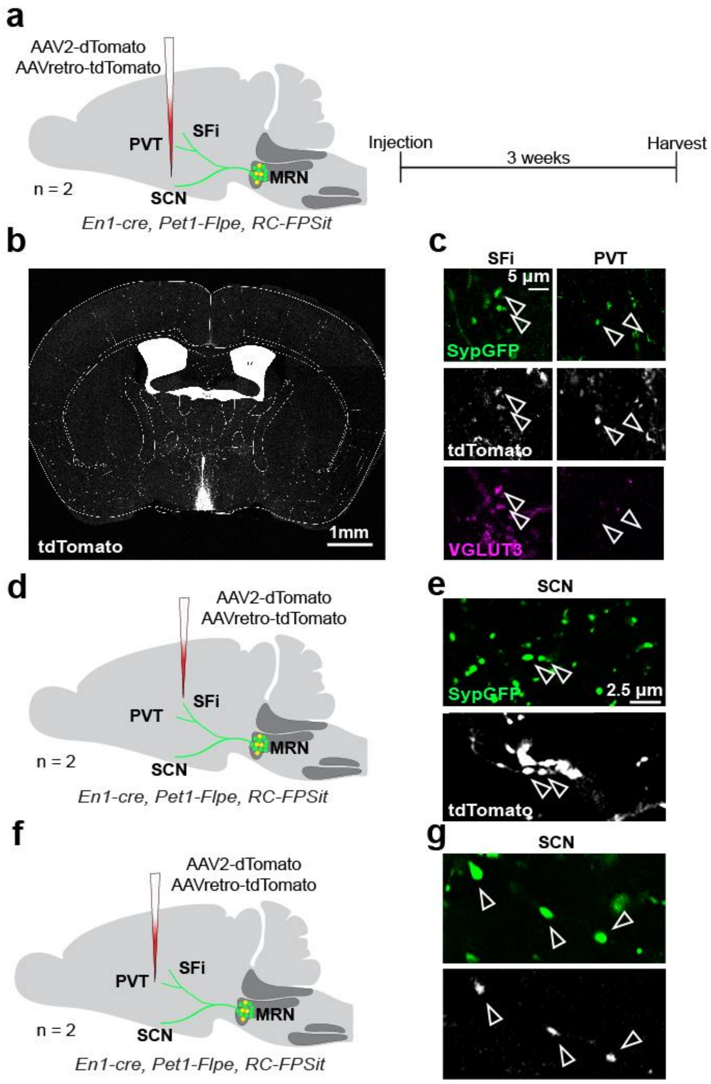


**Extended Data Fig. 3. SCN-projecting mr*En1-Pet1* neurons collateralize to PVT and SFi. a,** Schematic of the experimental approach and timeline. **b,** Representative image of the injection site. **c,** Representative images showing that collaterals in the SFi from SCN-projecting mr*En1-Pet1* neurons stain positive for VGLUT3, while collaterals in the PVT do not. n = 2. **d,** Schematic of the experimental approach. **e,** Representative image showing presence of SFi-projecting mr*En1-Pet1* neurons' collaterals in the SCN. n = 2. **f,** Schematic of the experimental approach. **g,** Representative image showing presence of PVT-projecting mr*En1-Pet1* neurons' collaterals in the SCN. n = 2.

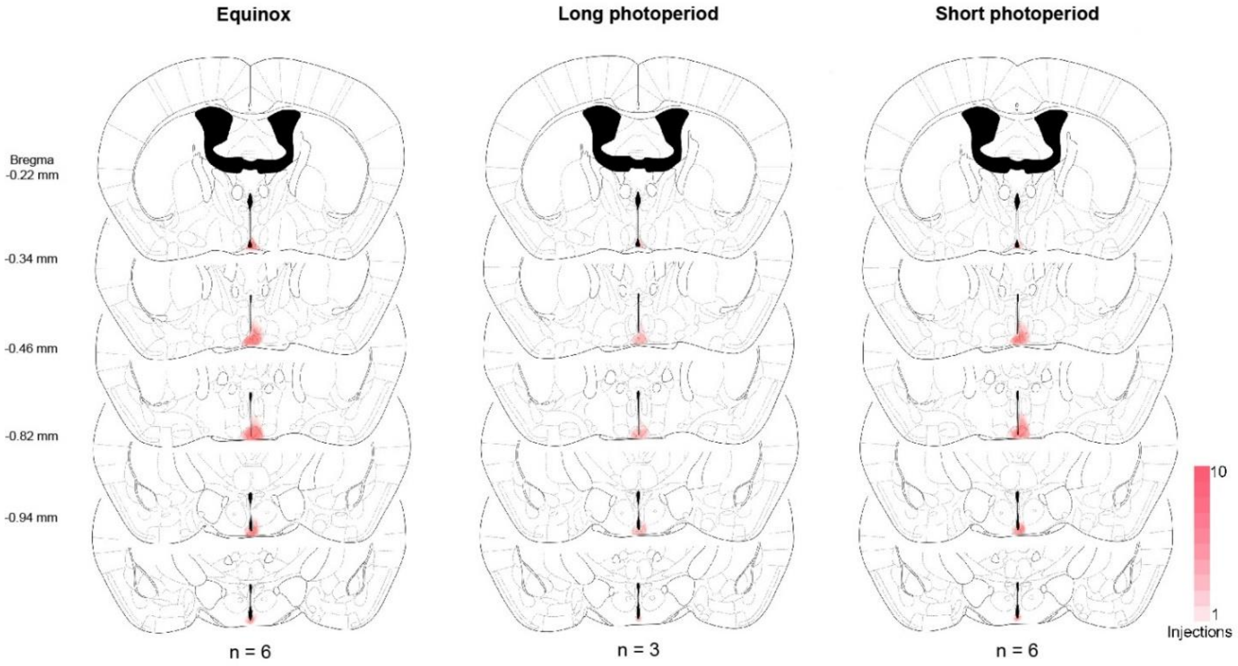


**Extended Data Fig. 4. Validation of retrograde viral circuit tracing from the SCN in different photoperiods**. Schematic of color-coded viral targeting and spread across 5 coronal planes.

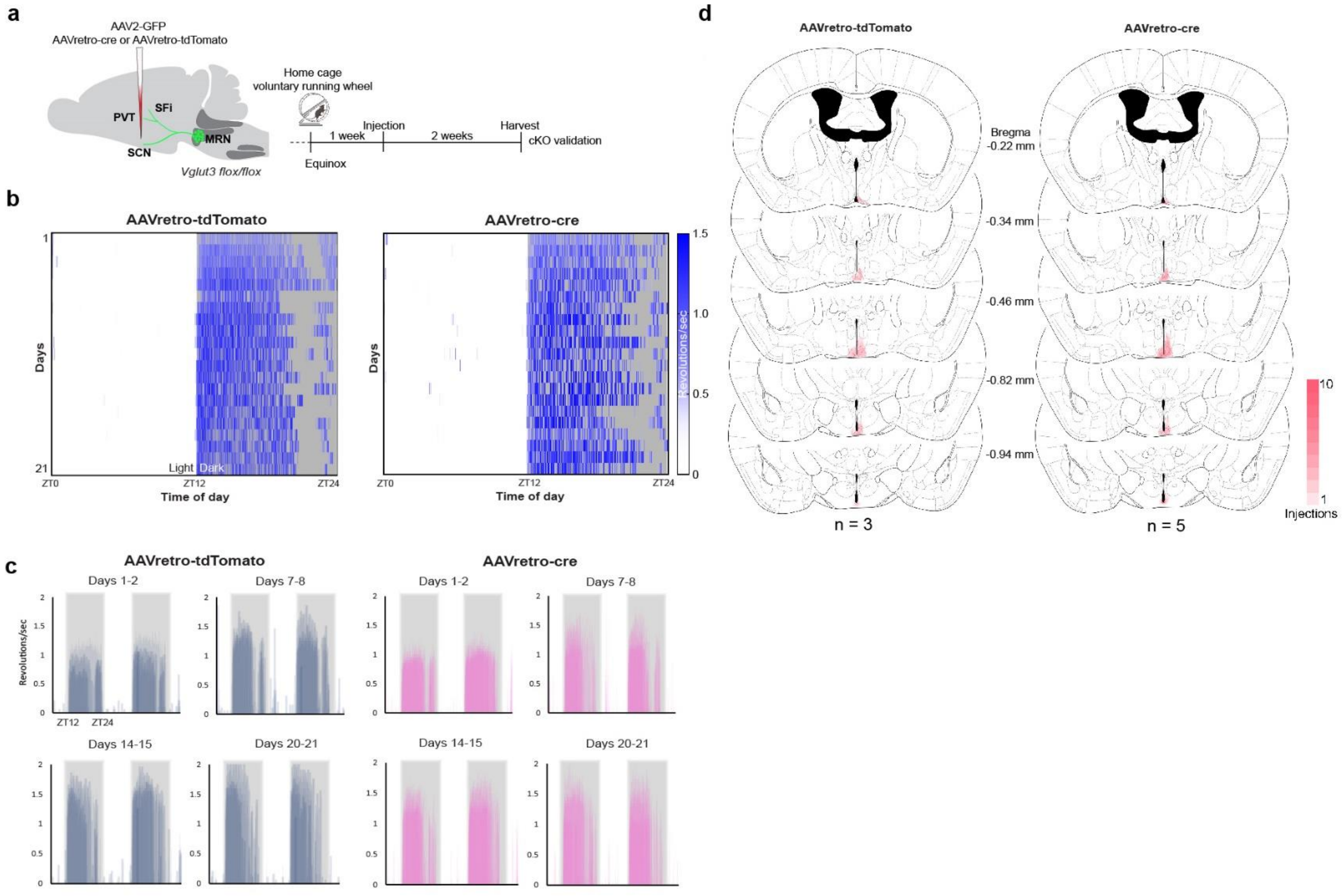


**Extended Data Fig. 5.** ***Vglut3*** **knockout in SCN-projecting serotonin neurons does not produce gross circadian abnormalities in equinox conditions. a,** Schematic of the experimental approach and timeline. **b,** Representative heatmaps of daily running activity (expressed as number of wheel revolutions per second) for control and cKO mice in equinox conditions. **c,** Wheel running activity during selected timepoints across the experimental timeline. **d,** Schematic of color-coded viral targeting and spread across 5 coronal planes.

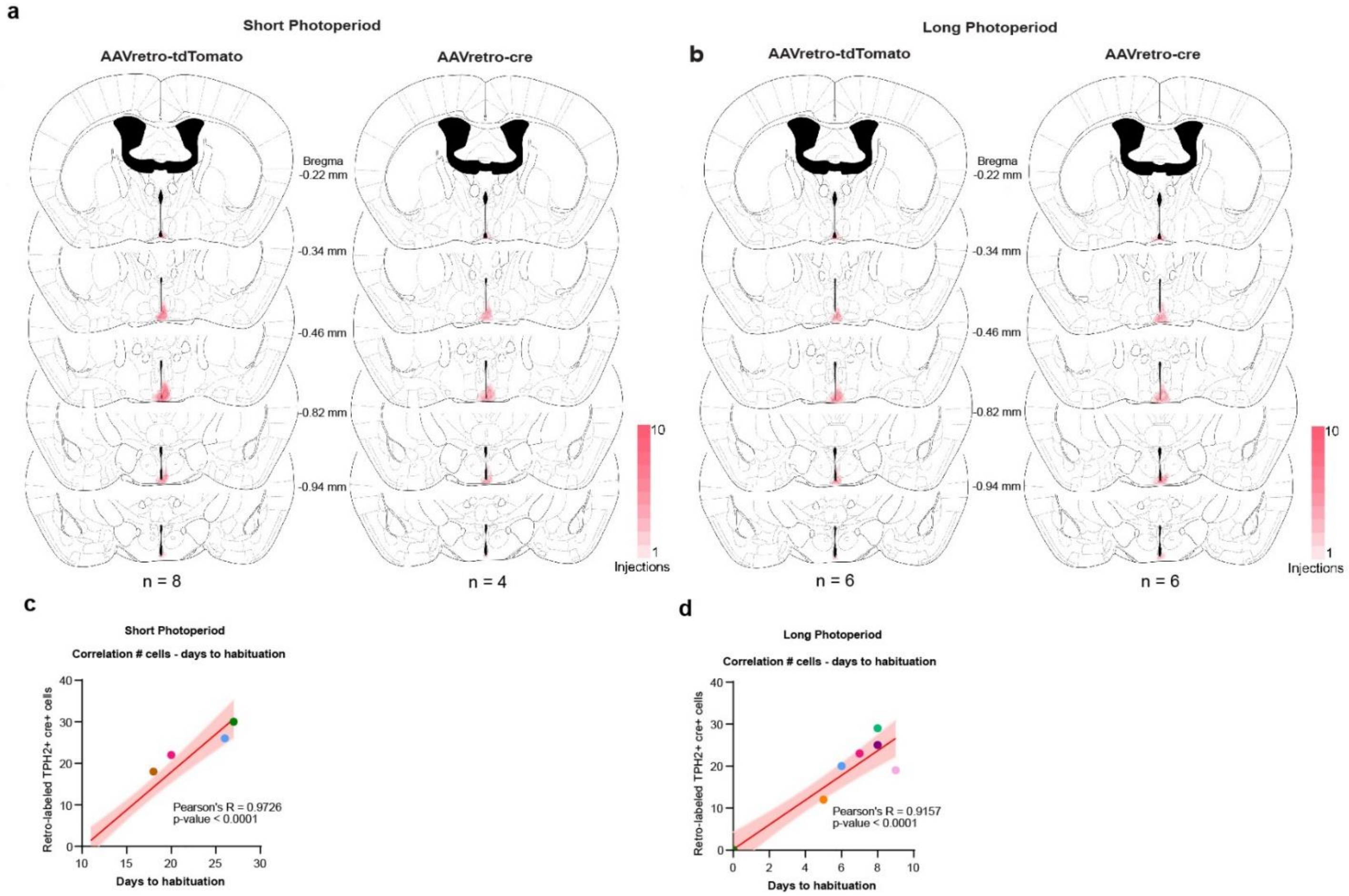


**Extended Data Fig. 6. Validation of viral targeting and correlation knock-out efficiency-running behaviour. a-b,** Schematic of color-coded viral targeting and spread across 5 coronal planes. **c,** Correlation between number of retrogradely labelled TPH2+ cells and days required to synchronize activity to short photoperiod changes. **d,** Correlation between number of retrogradely labelled TPH2+ cells and days required to synchronize activity to long photoperiod changes.

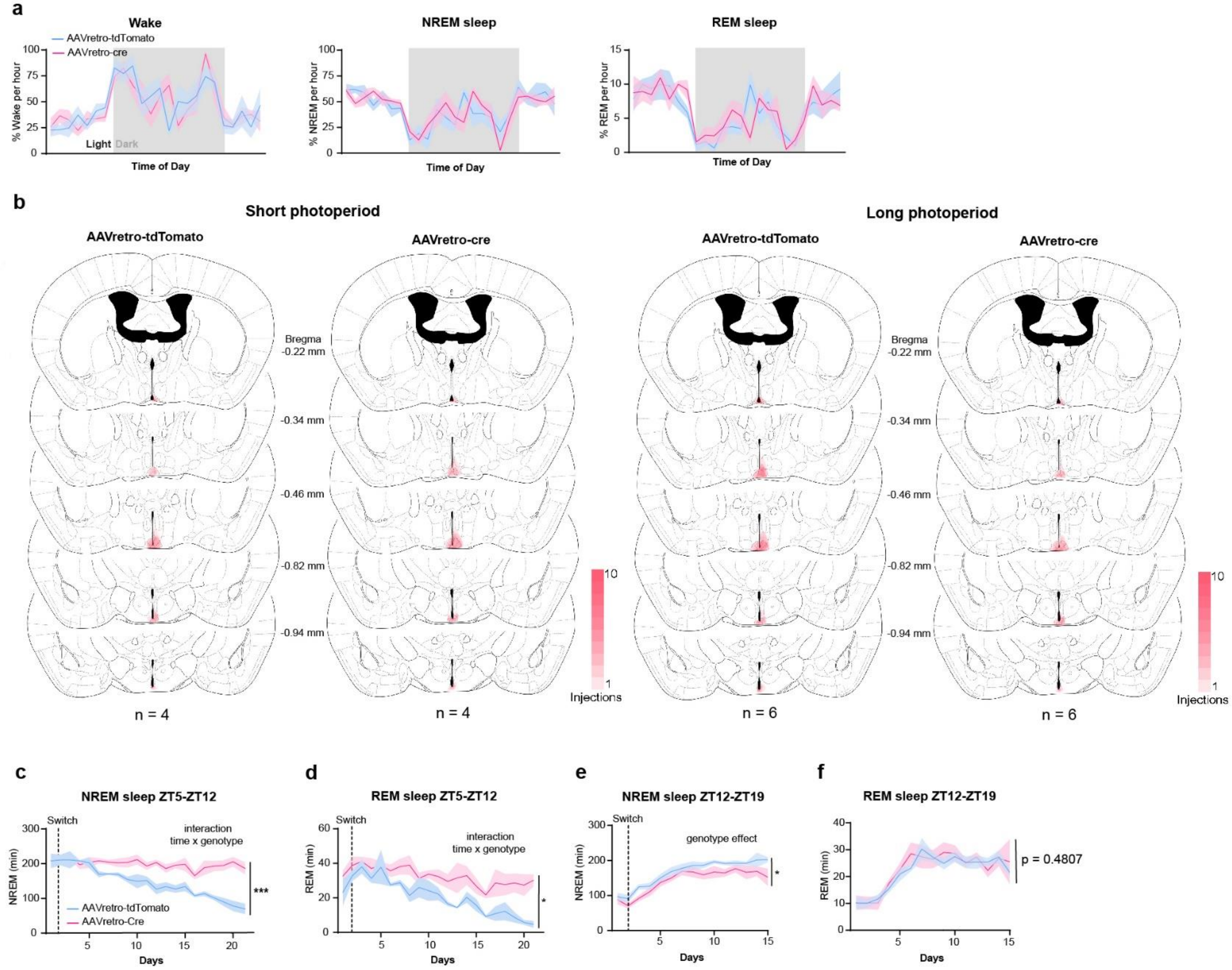


**Extended Data Fig. 7. *Vglut3* knockout in SCN-projecting serotonin neurons does not produce gross sleep/wake abnormalities in equinox conditions but disrupts sleep/wake adaptation to changes in daylength. a,** Time course analysis showing amount of Wake, NREM and REM sleep in equinox conditions. **b,** Schematic of color-coded viral targeting and spread across 5 coronal planes. **c,** Time spent in NREM sleep between ZT5 and ZT12 (Mixed-effect model, interaction genotype x time, $F$ (20, 54) = 11.19, p<0.0001, followed by Sidak's multiple comparison test). **d,** Time spent in REM sleep between ZT5 and ZT12 (Mixed-effect model, interaction genotype x time, $F$ (20, 53) = 2.238, p = 0.0101, followed by Sidak's multiple comparison test). **e,** Time spent in NREM sleep between ZT12 and ZT19 (two-way ANOVA, genotype effect, $F$ (1, 117) = 24.87, p<0.0001, followed by Sidak's multiple comparison test). **f,** Time spent in REM sleep between ZT12 and ZT19 (two-way ANOVA, genotype effect, $F$ (1, 117) = 20.5005, p = 0.4807, followed by Sidak's multiple comparison test).

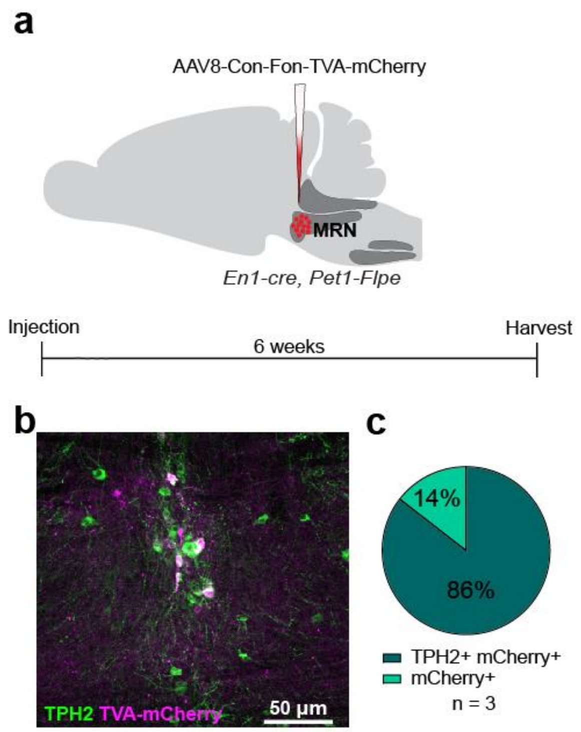


**Extended Data Fig. 8. Specificity of Cre- and Flp-dependent AAV encoding TVA-mCherry in mr*En1-Pet1* neurons. a,** Schematic of the experimental approach and experimental timeline. **b,** Representative image of transduced mr*En1-Pet1* neurons. **c,** Colocalization mCherry-TPH2. n=3 mice.

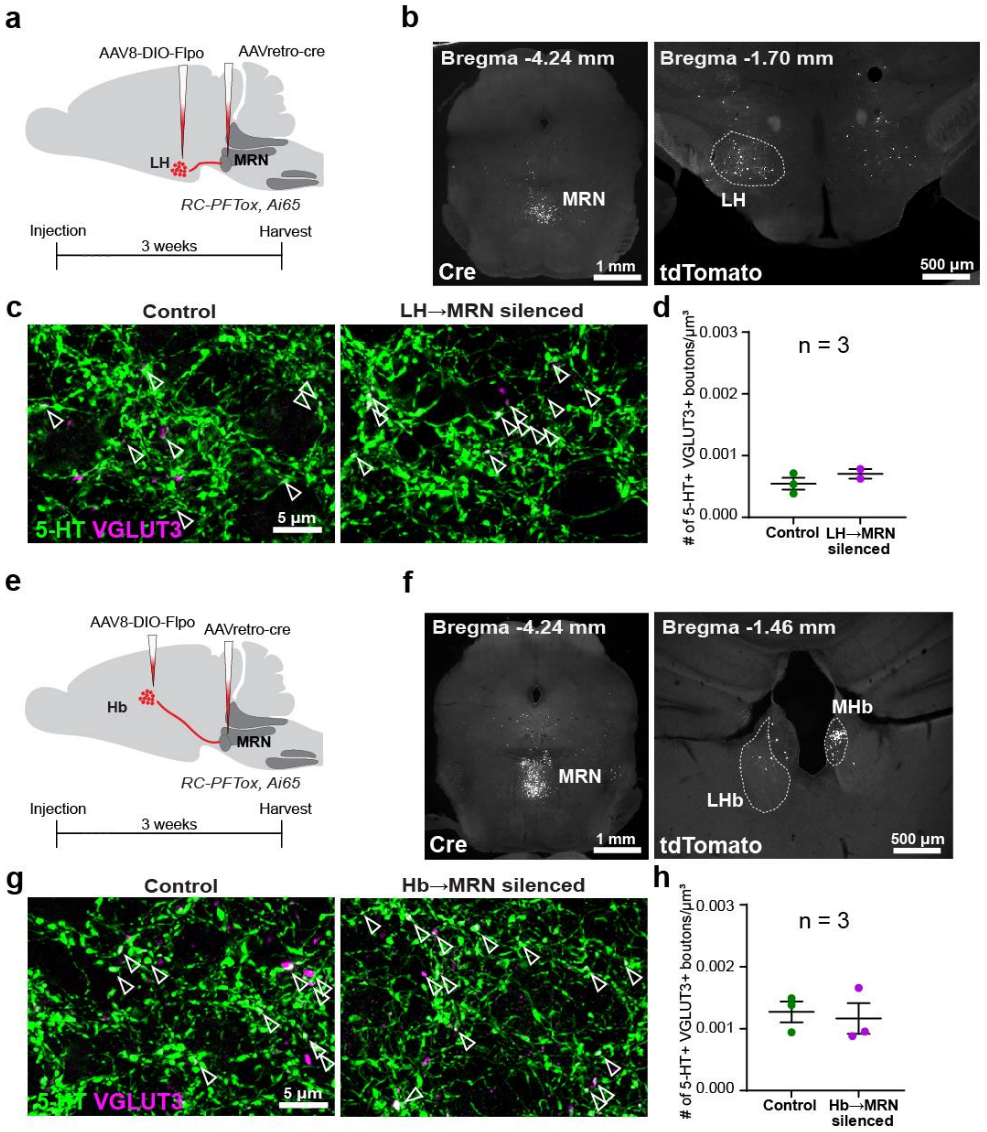


**Extended Data Fig. 9. Chronic silencing of LH→MRN and Hb→MRN does not promote 5-HT-VGLUT3 reorganization in mr*En1-Pet1*$^{\rightarrow SCN}$ boutons in the SCN. a,** Schematic of the experimental approach and experimental timeline. **b,** Representative image of the injection site in the MRN (right) and of the intersectionally labelled LH neuronal population (left). **c,** Representative superresolution images showing no difference in 5-HT and VGLUT3 overlap in the SCN of mice in which LH→MRN pathway has been chronically silenced. **d,** Quantification SCN 5-HT-VGLUT3 overlap in control and LH→MRN silenced mice. **e,** Schematic of the experimental approach and timeline. **f,** Representative image of the injection site in the MRN (right) and of the intersectionally labelled Hb neuronal population (left). **g,** Representative superresolution images showing no difference in 5-HT and VGLUT3 overlap in the SCN of mice in which Hb→MRN pathway has been chronically silenced. **h,** Quantification SCN 5-HT-VGLUT3 overlap in control and Hb→MRN silenced mice.

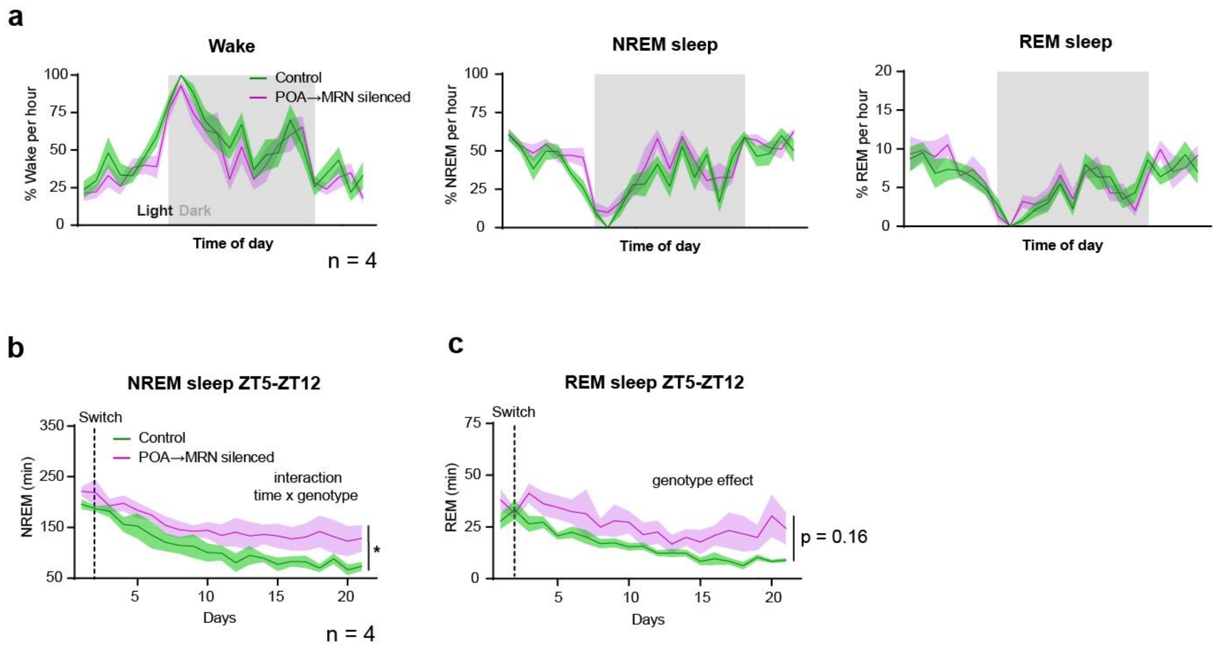


**Extended Data Fig. 10. Chronic silencing of POA→MRN neurons does not produce gross sleep/wake abnormalities in equinox conditions but disrupts sleep/wake adaptation to short photoperiod. a,** Time course analysis showing amount of Wake, NREM and REM sleep in equinox conditions. **b,** Time spent in NREM sleep between ZT5 and ZT12 (two-way ANOVA, interaction time x genotype, $F$ (20, 60) = 1.923, p = 0.0270, followed by Sidak's multiple comparison test). **c,** Time spent in REM sleep between ZT5 and ZT12 (two-way ANOVA, genotype effect, $F$ (20, 60) = 1.394, p = 0.1640, followed by Sidak's multiple comparison test).

### Multiple comparison test tables

#### Short photoperiod running wheel analyses

**Fig. 3f.** Activity onset. Sidak's test

| cKO vs control | Summary | Adjusted p-value |
|---|---|---|
| Day 1 | ns | >0.9999 |
| Day 2 | ns | >0.9999 |
| Day 3 | ns | >0.9999 |
| Day 4 | ns | >0.9999 |
| Day 5 | ns | >0.9999 |
| Day 6 | ns | >0.9999 |
| Day 7 | ns | >0.9999 |
| Day 8 | ns | >0.9999 |
| Day 9 | ns | >0.9999 |
| Day 10 | ns | >0.9999 |
| Day 11 | ns | 0.4829 |
| Day 12 | ns | 0.2068 |
| Day 13 | * | 0.0274 |
| Day 14 | ** | 0.0013 |
| Day 15 | ** | 0.0018 |
| Day 16 | *** | 0.0004 |
| Day 17 | *** | <0.0001 |
| Day 18 | ** | 0.0016 |
| Day 19 | *** | 0.0004 |
| Day 20 | *** | <0.0001 |
| Day 21 | *** | <0.0001 |
| Day 22 | * | 0.0211 |
| Day 23 | ns | 0.0857 |
| Day 24 | ns | 0.1148 |
| Day 25 | ns | 0.5001 |
| Day 26 | ns | 0.6512 |
| Day 27 | ns | 0.6537 |
| Day 28 | ns | 0.8522 |
| Day 29 | ns | 0.9949 |
| Day 30 | ns | >0.9999 |
| Day 31 | ns | >0.9999 |
| Day 32 | ns | >0.9999 |
| Day 33 | ns | >0.9999 |

**Fig. 3g.** Time spent running. Sidak's test

| cKO vs control | Summary | Adjusted p-value |
|---|---|---|
| Day -5 | ns | >0.9999 |
| Day -4 | ns | >0.9999 |
| Day -3 | ns | >0.9999 |
| Day -2 | ns | >0.9999 |
| Day -1 | ns | >0.9999 |
| Day 0 | ns | >0.9999 |
| Day 1 | ns | >0.9999 |
| Day 2 | ns | >0.9999 |
| Day 3 | ns | 0.9995 |
| Day 4 | ns | 0.5644 |
| Day 5 | ns | 0.9607 |
| Day 6 | ns | 0.4894 |
| Day 7 | ns | 0.2059 |
| Day 8 | * | 0.1304 |
| Day 9 | ns | 0.0335 |
| Day 10 | ** | 0.0529 |
| Day 11 | ns | 0.0023 |
| Day 12 | ns | 0.5760 |
| Day 13 | ns | 0.4595 |
| Day 14 | * | 0.3147 |
| Day 15 | * | 0.0442 |
| Day 16 | ns | 0.0162 |
| Day 17 | ns | 0.8779 |
| Day 18 | ns | 0.2672 |
| Day 19 | ns | 0.3609 |
| Day 20 | ns | 0.5230 |
| Day 21 | ns | >0.9999 |

**Long photoperiod running wheel analyses**

**Fig. 3i.** Activity onset. Sidak's test

| cKO vs control | Summary | Adjusted p-value |
|---|---|---|
| Day 1 | ns | >0.9999 |
| Day 2 | ns | >0.9999 |
| Day 3 | ns | >0.9999 |
| Day 4 | ns | >0.9999 |
| Day 5 | ns | >0.9999 |
| Day 6 | *** | <0.0001 |
| Day 7 | *** | <0.0001 |
| Day 8 | *** | <0.0001 |
| Day 9 | *** | <0.0001 |
| Day 10 | *** | <0.0001 |
| Day 11 | *** | <0.0001 |
| Day 12 | * | 0.0429 |
| Day 13 | ns | 0.1719 |
| Day 14 | ns | 0.9955 |
| Day 15 | ns | 0.8276 |
| Day 16 | ns | >0.9999 |

**Fig. 3j.** Time spent running. Sidak's test

| cKO vs control | Summary | Adjusted p-value |
|---|---|---|
| Day -5 | ns | >0.9999 |
| Day -4 | ns | >0.9999 |
| Day -3 | ns | >0.9999 |
| Day -2 | ns | >0.9999 |
| Day -1 | ns | >0.9999 |
| Day 0 | * | 0.0485 |
| Day 1 | ns | 0.0991 |
| Day 2 | ns | 0.5753 |
| Day 3 | ns | 0.8875 |
| Day 4 | ns | 0.8885 |
| Day 5 | ns | 0.9333 |
| Day 6 | ns | 0.9995 |
| Day 7 | ns | 0.9985 |

**Short photoperiod sleep analyses**

**Fig. 4c.** Wake ZT5-ZT12. Sidak's test.

| cKO vs control | Summary | Adjusted p-value |
|---|---|---|
| Day 1 | ns | >0.9999 |
| Day 2 | ns | >0.9999 |
| Day 3 | ns | >0.9999 |
| Day 4 | ns | >0.9999 |
| Day 5 | ns | 0.9993 |
| Day 6 | ns | 0.0899 |
| Day 7 | ns | 0.2365 |
| Day 8 | * | 0.0249 |
| Day 9 | ** | 0.0088 |
| Day 10 | *** | <0.0001 |
| Day 11 | ** | 0.0043 |
| Day 12 | *** | <0.0001 |
| Day 13 | *** | <0.0001 |
| Day 14 | *** | <0.0001 |
| Day 15 | *** | 0.0002 |
| Day 16 | *** | <0.0001 |
| Day 17 | *** | <0.0001 |
| Day 18 | *** | <0.0001 |
| Day 19 | *** | <0.0001 |
| Day 20 | *** | <0.0001 |
| Day 21 | *** | <0.0001 |

**Fig. 4d.** Sleep ZT5-ZT12. Sidak's test.

| cKO vs control | Summary | Adjusted p-value |
|---|---|---|
| Day 1 | ns | >0.9999 |
| Day 2 | ns | >0.9999 |
| Day 3 | ns | >0.9999 |
| Day 4 | ns | >0.9999 |
| Day 5 | ns | >0.9999 |
| Day 6 | ns | 0.2197 |
| Day 7 | ns | 0.4945 |
| Day 8 | ns | 0.1733 |
| Day 9 | ns | 0.1054 |
| Day 10 | ** | 0.0011 |
| Day 11 | ns | 0.0717 |
| Day 12 | *** | 0.0009 |
| Day 13 | ** | 0.0070 |
| Day 14 | ** | 0.0013 |
| Day 15 | ** | 0.0080 |
| Day 16 | ** | 0.0014 |
| Day 17 | *** | 0.0001 |
| Day 18 | *** | <0.0001 |
| Day 19 | *** | <0.0001 |
| Day 20 | *** | <0.0001 |
| Day 21 | *** | <0.0001 |

**Extended Fig. 7c.** NREM ZT5-ZT12. Sidak's test.

| cKO vs control | Summary | Adjusted p-value |
|---|---|---|
| Day 1 | ns | >0.9999 |
| Day 2 | ns | >0.9999 |
| Day 3 | ns | >0.9999 |
| Day 4 | ns | >0.9999 |
| Day 5 | ns | >0.9999 |
| Day 6 | ns | 0.1661 |
| Day 7 | ns | 0.5230 |
| Day 8 | ns | 0.4506 |
| Day 9 | * | 0.0322 |
| Day 10 | ** | 0.0014 |
| Day 11 | ns | 0.0694 |
| Day 12 | *** | 0.0003 |
| Day 13 | * | 0.0124 |
| Day 14 | *** | 0.0004 |
| Day 15 | ** | 0.0046 |
| Day 16 | ** | 0.0067 |
| Day 17 | *** | 0.0001 |
| Day 18 | *** | <0.0001 |
| Day 19 | *** | <0.0001 |
| Day 20 | *** | <0.0001 |
| Day 21 | *** | <0.0001 |

**Extended Fig. 7d.** REM ZT5-ZT12. Sidak's test.

| cKO vs control | Summary | Adjusted p-value |
|---|---|---|
| Day 1 | ns | 0.6151 |
| Day 2 | ns | 0.9994 |
| Day 3 | ns | >0.9999 |
| Day 4 | ns | >0.9999 |
| Day 5 | ns | >0.9999 |
| Day 6 | ns | 0.9762 |
| Day 7 | ns | 0.9403 |
| Day 8 | ns | 0.0708 |
| Day 9 | ns | >0.9999 |
| Day 10 | ns | 0.5114 |
| Day 11 | ns | 0.6659 |
| Day 12 | ns | 0.6725 |
| Day 13 | ns | 0.0613 |
| Day 14 | ns | 0.5869 |
| Day 15 | ns | 0.7222 |
| Day 16 | ns | 0.4902 |
| Day 17 | * | 0.0253 |
| Day 18 | ns | 0.0785 |
| Day 19 | * | 0.0101 |
| Day 20 | ** | 0.0046 |
| Day 21 | *** | 0.0005 |

**Long photoperiod sleep analyses**

**Fig. 4f.** Wake ZT12-ZT19. Sidak's test.

| cKO vs control | Summary | Adjusted p-value |
|---|---|---|
| Day 1 | ns | >0.9999 |
| Day 2 | ns | 0.9918 |
| Day 3 | ns | 0.2036 |
| Day 4 | ns | 0.9740 |
| Day 5 | ns | 0.8748 |
| Day 6 | ns | 0.8129 |
| Day 7 | ns | 0.6276 |
| Day 8 | ns | 0.5305 |
| Day 9 | ns | 0.1793 |
| Day 10 | ns | 0.1381 |
| Day 11 | * | 0.0499 |
| Day 12 | ns | 0.4786 |
| Day 13 | ns | 0.1077 |
| Day 14 | * | 0.0149 |

**Fig. 4g.** Sleep ZT12-ZT19. Sidak's test.

| cKO vs control | Summary | Adjusted p-value |
|---|---|---|
| Day 1 | ns | 0.9990 |
| Day 2 | ns | >0.9999 |
| Day 3 | ns | 0.5289 |
| Day 4 | ns | >0.9999 |
| Day 5 | ns | 0.9996 |
| Day 6 | ns | >0.9999 |
| Day 7 | ns | >0.9999 |
| Day 8 | ns | 0.9483 |
| Day 9 | ns | 0.8863 |
| Day 10 | ns | 0.5253 |
| Day 11 | ns | 0.9808 |
| Day 12 | ns | 0.9016 |
| Day 13 | ns | 0.7875 |
| Day 14 | * | 0.0498 |

**Extended Fig. 7e.** NREM ZT12-ZT19. Sidak's test.

| cKO vs control | Summary | Adjusted p-value |
|---|---|---|
| Day 1 | ns | >0.9999 |
| Day 2 | ns | 0.9770 |
| Day 3 | ns | 0.6332 |
| Day 4 | ns | 0.9985 |
| Day 5 | ns | >0.9999 |
| Day 6 | ns | 0.9908 |
| Day 7 | ns | >0.9999 |
| Day 8 | ns | 0.9917 |
| Day 9 | ns | 0.7970 |
| Day 10 | ns | 0.6967 |
| Day 11 | ns | 0.8953 |
| Day 12 | ns | 0.9995 |
| Day 13 | ns | 0.9288 |
| Day 14 | ns | 0.7885 |

**Extended Fig. 7f.** REM ZT12-ZT19. Sidak's test.

| cKO vs control | Summary | Adjusted p-value |
|---|---|---|
| Day 1 | ns | >0.9999 |
| Day 2 | ns | >0.9999 |
| Day 3 | ns | >0.9999 |
| Day 4 | ns | >0.9999 |
| Day 5 | ns | >0.9999 |
| Day 6 | ns | 0.9142 |
| Day 7 | ns | 0.9998 |
| Day 8 | ns | >0.9999 |
| Day 9 | ns | 0.9961 |
| Day 10 | ns | >0.9999 |
| Day 11 | ns | >0.9999 |
| Day 12 | ns | >0.9999 |
| Day 13 | ns | 0.9999 |
| Day 14 | ns | >0.9999 |

**Short photoperiod sleep analyses POA-MRN**

**Fig. 6g.** Wake ZT5-ZT12. Sidak's test.

| cKO vs control | Summary | Adjusted p-value |
|---|---|---|
| Day 1 | ns | 0.7164 |
| Day 2 | ns | 0.4738 |
| Day 3 | * | 0.0329 |
| Day 4 | *** | <0.0001 |
| Day 5 | ** | 0.0023 |
| Day 6 | *** | <0.0001 |
| Day 7 | ** | 0.0028 |
| Day 8 | * | 0.0224 |
| Day 9 | ** | 0.0092 |
| Day 10 | *** | 0.0010 |
| Day 11 | ** | 0.0044 |
| Day 12 | *** | <0.0001 |
| Day 13 | ** | 0.0041 |
| Day 14 | *** | 0.0001 |
| Day 15 | *** | <0.0001 |
| Day 16 | *** | 0.0002 |
| Day 17 | *** | <0.0001 |
| Day 18 | *** | <0.0001 |
| Day 19 | *** | <0.0001 |
| Day 20 | *** | <0.0001 |
| Day 21 | *** | <0.0001 |

**Fig. 6g.** Sleep ZT5-ZT12. Sidak's test.

| cKO vs control | Summary | Adjusted p-value |
|---|---|---|
| Day 1 | ns | 0.2262 |
| Day 2 | ns | 0.9999 |
| Day 3 | ns | 0.7716 |
| Day 4 | ns | 0.7729 |
| Day 5 | * | 0.0381 |
| Day 6 | * | 0.0147 |
| Day 7 | * | 0.0431 |
| Day 8 | ns | 0.1270 |
| Day 9 | ns | 0.0815 |
| Day 10 | ** | 0.0031 |
| Day 11 | ns | 0.0962 |
| Day 12 | *** | <0.0001 |
| Day 13 | ns | 0.0613 |
| Day 14 | ** | 0.0047 |
| Day 15 | *** | 0.0003 |
| Day 16 | ** | 0.0032 |
| Day 17 | *** | 0.0004 |
| Day 18 | *** | <0.0001 |
| Day 19 | ** | 0.0012 |
| Day 20 | *** | <0.0001 |
| Day 21 | *** | <0.0001 |

**Extended Fig. 10b.** NREM ZT5-ZT12. Sidak's test.

| cKO vs control | Summary | Adjusted p-value |
|---|---|---|
| Day 1 | ns | 0.2455 |
| Day 2 | * | 0.0494 |
| Day 3 | ns | 0.9990 |
| Day 4 | ** | 0.0020 |
| Day 5 | ns | 0.0570 |
| Day 6 | ** | 0.0044 |
| Day 7 | * | 0.0311 |
| Day 8 | ns | 0.0587 |
| Day 9 | ns | 0.0853 |
| Day 10 | *** | 0.0009 |
| Day 11 | * | 0.0191 |
| Day 12 | *** | <0.0001 |
| Day 13 | ** | 0.0060 |
| Day 14 | *** | 0.0003 |
| Day 15 | *** | <0.0001 |
| Day 16 | *** | 0.0007 |
| Day 17 | *** | 0.0001 |
| Day 18 | *** | <0.0001 |
| Day 19 | ** | 0.0011 |
| Day 20 | *** | <0.0001 |
| Day 21 | *** | <0.0001 |

**Extended Fig. 10c.** REM ZT5-ZT12. Sidak's test.

| cKO vs control | Summary | Adjusted p-value |
|---|---|---|
| Day 1 | ns | 0.2392 |
| Day 2 | ns | >0.9999 |
| Day 3 | * | 0.0118 |
| Day 4 | ns | 0.4895 |
| Day 5 | * | 0.0238 |
| Day 6 | ns | 0.2933 |
| Day 7 | ns | 0.1460 |
| Day 8 | ns | 0.6805 |
| Day 9 | ns | 0.1986 |
| Day 10 | ns | 0.0926 |
| Day 11 | ns | 0.9784 |
| Day 12 | ns | 0.2624 |
| Day 13 | ns | 0.9993 |
| Day 14 | ns | 0.7381 |
| Day 15 | ns | 0.3928 |
| Day 16 | ns | 0.1462 |
| Day 17 | * | 0.0107 |
| Day 18 | ** | 0.0070 |
| Day 19 | ns | 0.3520 |
| Day 20 | *** | <0.0001 |
| Day 21 | ** | 0.0082 |
